# BingleSeq: A user-friendly R package for Bulk and Single-cell RNA-Seq Data Analysis

**DOI:** 10.1101/2020.06.16.148239

**Authors:** Daniel Dimitrov, Quan Gu

## Abstract

RNA sequencing is a high-throughput sequencing technique considered as an indispensable research tool used in a broad range of transcriptome analysis studies. The most common application of RNA Sequencing is Differential Expression analysis and it is used to determine genetic loci with distinct expression across different conditions. On the other hand, an emerging field called single-cell RNA sequencing is used for transcriptome profiling at the individual cell level. The standard protocols for both these types of analyses include the processing of sequencing libraries and result in the generation of count matrices. An obstacle to these analyses and the acquisition of meaningful results is that both require programming expertise.

BingleSeq was developed as an intuitive application that provides a user-friendly solution for the analysis of count matrices produced by both Bulk and Single-cell RNA-Seq experiments. This was achieved by building an interactive dashboard-like user interface and incorporating three state-of-the-art software packages for each type of the aforementioned analyses, alongside additional features such as key visualisation techniques, functional gene annotation analysis and rank-based consensus for differential gene analysis results, among others. As a result, BingleSeq puts the best and most widely used packages and tools for RNA-Seq analyses at the fingertips of biologists with no programming experience.

## Introduction

About a decade ago, a transcriptome profiling approach, known as RNA Sequencing (RNA-Seq), was predicted to revolutionize transcriptome analyses (1). Today, as a consequence of the continuous advancements and dropping costs of next-generation sequencing (NGS) technologies, differential expression (DE) analysis or Bulk RNA-Seq has established itself as a routine research tool (2).

Single-cell RNA Sequencing (scRNA-Seq) is an emerging field that enables the gene expression profiling at the individual cell level. scRNA-Seq analysis is believed to lead to the reconstruction of an entire human cell lineage tree (3) and its potential is highlighted by its broad range of applications which include the classification and profiling of subpopulations of cells to putative transcriptomic types (4–6), the discovery of novel cell types (7,8) and novel microbial species (9), as well as the deconvolution of Bulk RNA-Seq results (10,11).

Although there is a wide range of software tools available for both Bulk RNA-Seq and scRNA-Seq analyses, most require some proficiency in programming languages such as R. This creates a challenge for the analysis of RNA-seq data for a large portion of biologists lacking programming experience. Here we present an application, called BingleSeq, the primary goal of which is to enable the user-friendly analysis of count tables obtained by both Bulk RNA-Seq and scRNA-Seq protocols.

## Design and Implementation

### Implementation

BingleSeq is based on shiny (12) and it is composed of a multi-tabbed UI, built as separate shiny modules with efficiency, code readability and reusability in mind. Each module (tab) corresponds to a key step in the typical Bulk and scRNA-Seq analysis pipelines. Modules are generated only upon reaching a given step of the analyses, which ensures efficiency and speed despite the complexity of the application. BingleSeq’s UI components (e.g. plots, tables, tabs, etc.) make use of shiny’s ‘reactivity’ property and these components are automatically updated upon user input or any change related to a given component. The application also enables users to customize key parameters for each step of the analyses according to the experiment in question, while also aiming to maintain an appropriate level of complexity. Consequently, BingleSeq provides an intuitive user experience, complemented with customizable plots and interactive tables as well as analysis-related tips and pop-up messages as further guidance to ensure the correct execution of the pipelines.

### Methods and Features

In terms of functionality, BingleSeq can be divided into two main parts corresponding to Bulk RNA-Seq and scRNA-Seq analysis pipelines (see **Fig. 1**).

**Figure 1.**
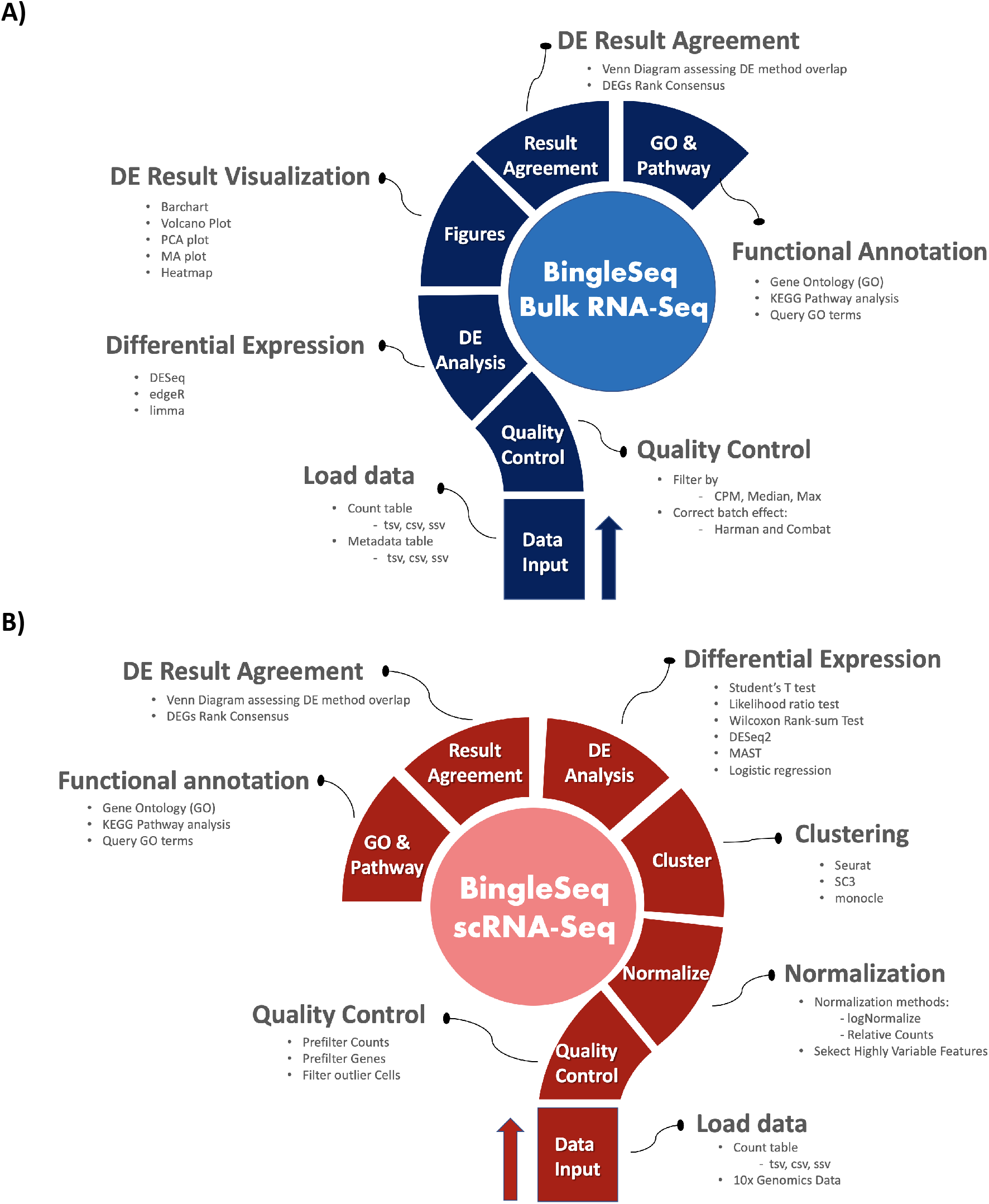
Overview of BingleSeq’s Bulk RNA-Seq (**A**) and scRNA-Seq (**B**) analysis pipelines.

As solutions to Bulk RNA-Seq analysis, BingleSeq implements DESeq2 (13), edgeR (14), and limma (15). These packages are well-tested and regarded as being among the best performing ones (16–18). Despite being accepted as being possibly the best DE analysis solutions, different studies often present contrasting conclusions. Hence, there is little consensus regarding which DE algorithm has the best performance. This stems from fact that there is no optimal package under all circumstances and different variables are known to affect package performance, with sample size in particular (16,18). Thus, the method of choice depends on the dataset being analyzed. The scRNA-Seq pipeline, includes three unsupervised clustering solutions provided by monocle (19), Seurat (20), and SC3 (21) packages. The latter two packages are regarded as having the best overall performance (22,23). However, similarly to packages used in the DE analysis of Bulk RNA-Seq data, there seems to be little consensus on which package provides the best-performing clustering approach. This is largely due to the inherent limitations of the different algorithms used in clustering, as a result no algorithm performs well in every circumstance (24). Kiselev et al. (25) suggest that Seurat may be inappropriate for small scRNA-Seq datasets, due to the inherent limitations of the Louvain algorithm. On the contrary, as a way to amend for the limitations of k-means clustering algorithm used in SC3 (21), the authors implemented an extensive iterative-consensus approach, which makes SC3 magnitudes slower than Seurat and downgrades its scalability (22,25). Another difference between these two packages is that Seurat does not include functionality to estimate or explicitly specify cluster number, while SC3 does.

In addition to enabling users to choose the most suitable package for their experiment, BingleSeq implements essential features such as quality control, batch effect correction, normalization, and a wide range of visualization techniques. The application also equips users with a solution for the extensive functional analysis of genes via the GOseq pipeline (26) and ‘GO.db’ package (27). The combination of these packages enables KEGG pathway and Gene ontology (GO) analyses as well as the retrieval of additional information about GO terms of interest. Furthermore, BingleSeq provides further confidence in DGEs and in the agreement between the methods using a rank-based consensus, complemented with a Venn Diagram.

### Related Applications

Some effort has been directed towards lowering the entry requirements to RNA-Seq analyses as there are some software tools which implement UI components. However, many of these applications are limited to only some key features or particular parts of RNA-Seq analysis (21,28). From the available software packages providing comprehensive solutions for Bulk RNA-Seq, DEapp (29), DEBrowser (30), and Omics Playground (31) were thought to provide the most options, with the latter two being more extensive in terms of additional features than DEapp. As seen in **S1-2 Table**, the functionality implemented BingleSeq’s Bulk RNA-Seq can be argued to put our package on par with even the best available Bulk RNA-Seq solutions.

When looking at similar applications providing solutions to scRNA-Seq analysis, BingleSeq is most analogous to SeuratWizard (32) as both are based on Seurat’s pipeline. However, by implementing SC3 and monocle, BingleSeq provides solutions to some of Seurat’s inherent limitations. For instance, SeuratWizard does not implement functionality to explicitly specify the number of clusters, nor a way to estimate the number of clusters, while BingleSeq provides two distinct approaches to achieve that. Another major functionality that sets our application apart is that it enables functional annotation analysis and a way to compare and provide a consensus for Seurat’s inbuilt DE methods. Consequently, BingleSeq can be argued to match even the most comprehensive scRNA-Seq applications, such as ASAP (33) and singleCellTK (34) – see **S2-3 Table**.

In terms of providing a solution to both Bulk and single-cell RNA-Seq analyses, BingleSeq’s features and comprehensiveness are only contested by those of Omics Playground. What sets BingleSeq apart is that it provides multiple clustering packages and algorithms as well as a larger number of biomarker visualization options.

## Results

### Bulk RNA-Seq Steps and Features

To begin the DE analysis of Bulk RNA-Seq data, a count table and a metadata table must be loaded in the appropriate formats (**S1 Fig**). The genes can then be filtered according to counts per million (CPM), Max, or Median thresholds and batch effect correction can be performed with Harman and ComBat packages (35,36) (**S2 Fig**).

Subsequent to Quality Control, users can investigate the differentially expressed genes (DEGs) using three state-of-the-art packages: DESeq2 (13), edgeR (14), and limma (15). To assess BingleSeq’s Bulk RNA-Seq pipeline, we used a synthetic dataset generated with compcodeR package (37) as well as a real data set looking at DEGs between HSV-1 infected control and interferon B treatment (38) – see **S1 File**.

Upon obtaining DE results, users can visualize them using key plotting techniques (**Fig. 2A-E**). These include a PCA plot used to provide insights about the relationship between samples; Barchart plot supplemented with a summary table which serve as an excellent way to summarize the up- and downregulated DEGs; Volcano and MA plots which are essential when assessing the relationship between fold change (FC) versus significance and average expression; and Heatmaps as they are thought to be the most versatile and informative type of visualization technique when looking at DE results.

**Figure 2.**
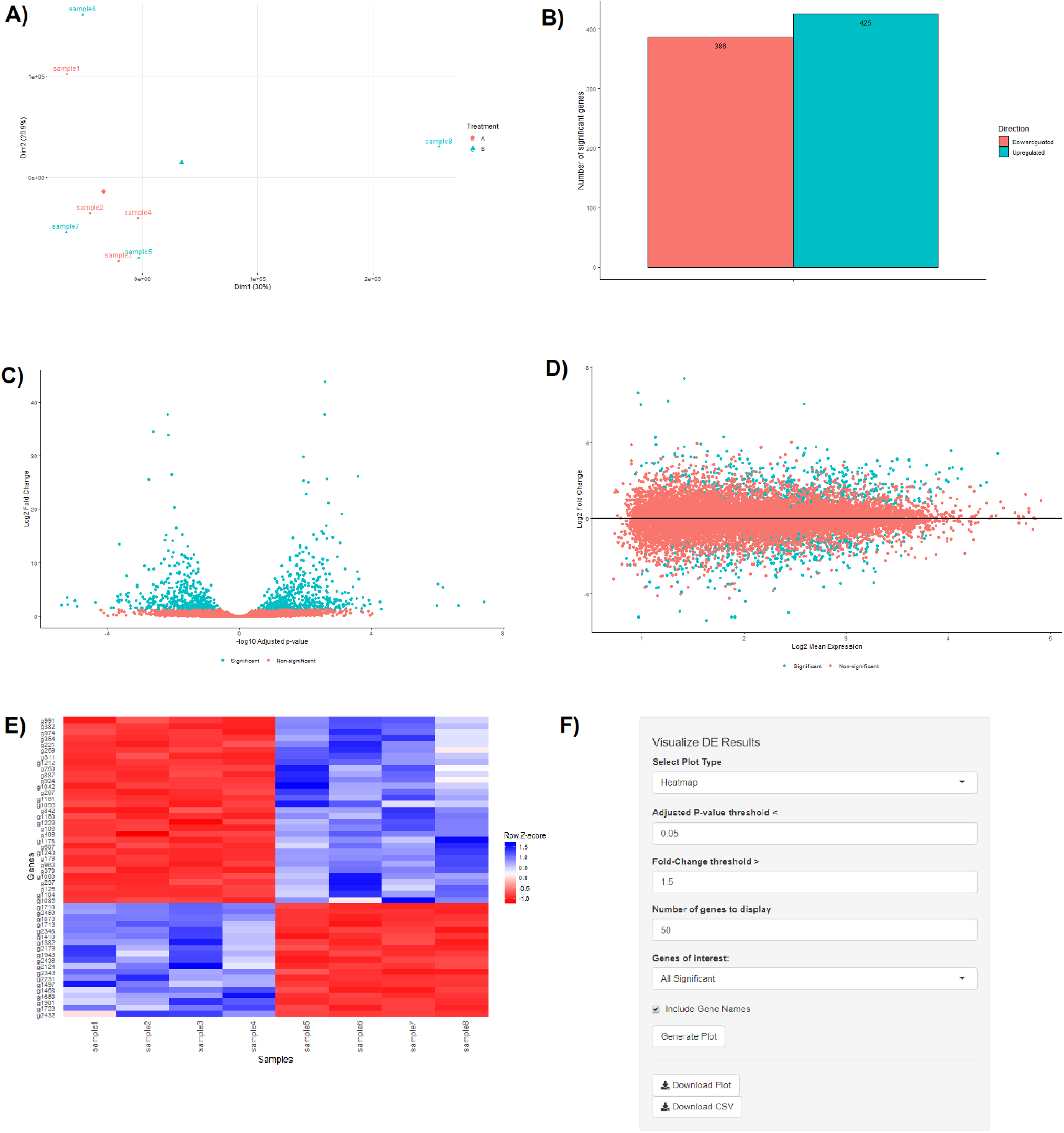
PCA plot (A), Barchart (B), Volcano plot (C), MA plot (D), and a Heatmap (E) with its corresponding interactive control panel (F) as generated by BingleSeq.

BingleSeq’s visualization techniques were implemented with customization in mind, as users can specify parameters such as p-value threshold and fold-change threshold, among others. Due to their versatility, heatmaps were designed as BingleSeq’s most customizable plotting component (**Fig. 2F**).

### scRNA-Seq Pipeline Steps and Features

The scRNA-Seq part is based on Seurat’s scRNA-Seq pipeline and visualization options (20). Nonetheless, clustering can also be performed with monocle (19) and SC3 (21) packages. An evaluation of BingleSeq’s scRNA-Seq pipeline was performed by reproducing and extending the results of Seurat’s online tutorial (https://satijalab.org/seurat/v3.0/pbmc3k_tutorial.html) - see **S2 File**. The tutorial is based on a 10x Genomics dataset of 2700 Peripheral Blood Mononuclear Cells (PBMCs) with ~69,000 reads per cell. The data set is available at https://support.10xgenomics.com/single-cell-gene-expression/datasets/1.1.0/pbmc3k.

To begin scRNA-Seq analysis, data can be supplied in two formats - ‘10x Genomics data’ as well as a count table with a specific format (**S3 Fig**). Once the data is loaded, users can filter unwanted cells and features, supplemented with visual aid in the form of violin plots (**S4 Fig**). Following filtering, the next step is to normalize the data and BingleSeq provides two Seurat-supplied normalization methods – “LogNormalize” and “Relative counts”. Simultaneously with normalization and scaling, the highly variable features within the dataset are identified and selected for clustering with Seurat (**S4 Fig**) as a way to minimize noise.

Following normalization, the ‘Clustering’ tab is generated (**Fig. 3**) which provides a high degree of control over the different steps of the analysis (**Fig. 3A**), alongside general tips for each step of the pipelines, such as clustering advice provided for each clustering package (**Fig. 3B**). The clustering tab is first used to perform pre-clustering prerequisites such as scaling the data and dimensionality reduction with Principal Component Analysis (PCA). Once these actions are performed, an elbow plot is returned which is used to determine the dimensionality of the dataset (**Fig. 3C**) – this is essential for excluding noise when clustering with Seurat and monocle. Also, PC heatmaps (**Fig. 3D**) are available as a further tool for PC Selection.

**Figure 3.**
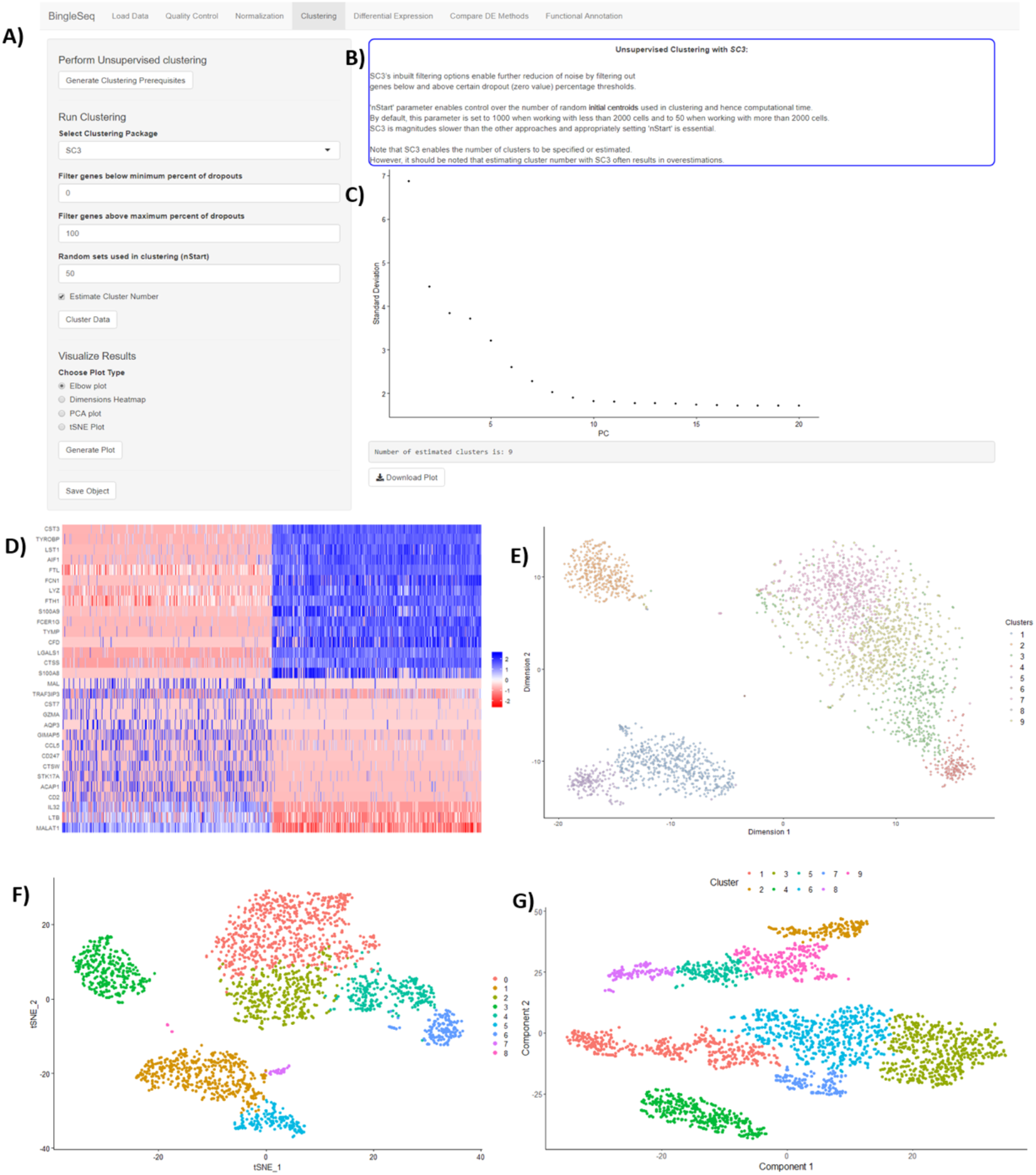
An overview of BingleSeq’s ‘Clustering’ tab with **A)** clustering customization options, **B)** general tips and advice for the selected clustering package (in this case SC3), and clustering-related visualization options: **C)** A PC elbow plot, **D)** a PC heatmap showing top 10 most variable genes in PC1, and tSNE plots produced with **E)** SC3, **F)** Seurat, and **G)** monocle.

Once the count data is filtered and transformed, users can proceed to unsupervised clustering with Seurat, SC3, and monocle. The primary way to visualize clustering results is via t-distributed stochastic neighbour embedding (tSNE) plots (**Fig. 3E-G**) – a method designed for the purpose of visualizing high dimensional datasets (39).

Following clustering, DE analysis can be conducted using Seurat’s inbuilt testing methods to identify marker genes. The implemented Seurat DE methods include: Student’s T test, Wilcoxon Rank Sum test, DESEq2 (13), and MAST package (42). DE results and specific marker genes can then be examined and visualized using Seurat’s inbuilt DE visualization options. These include cluster heatmap with user-specified gene number as well as visualizations for the exploration of specific genes via Violin, Feature, and Ridge plots (**Fig. 4**).

**Figure 4.**
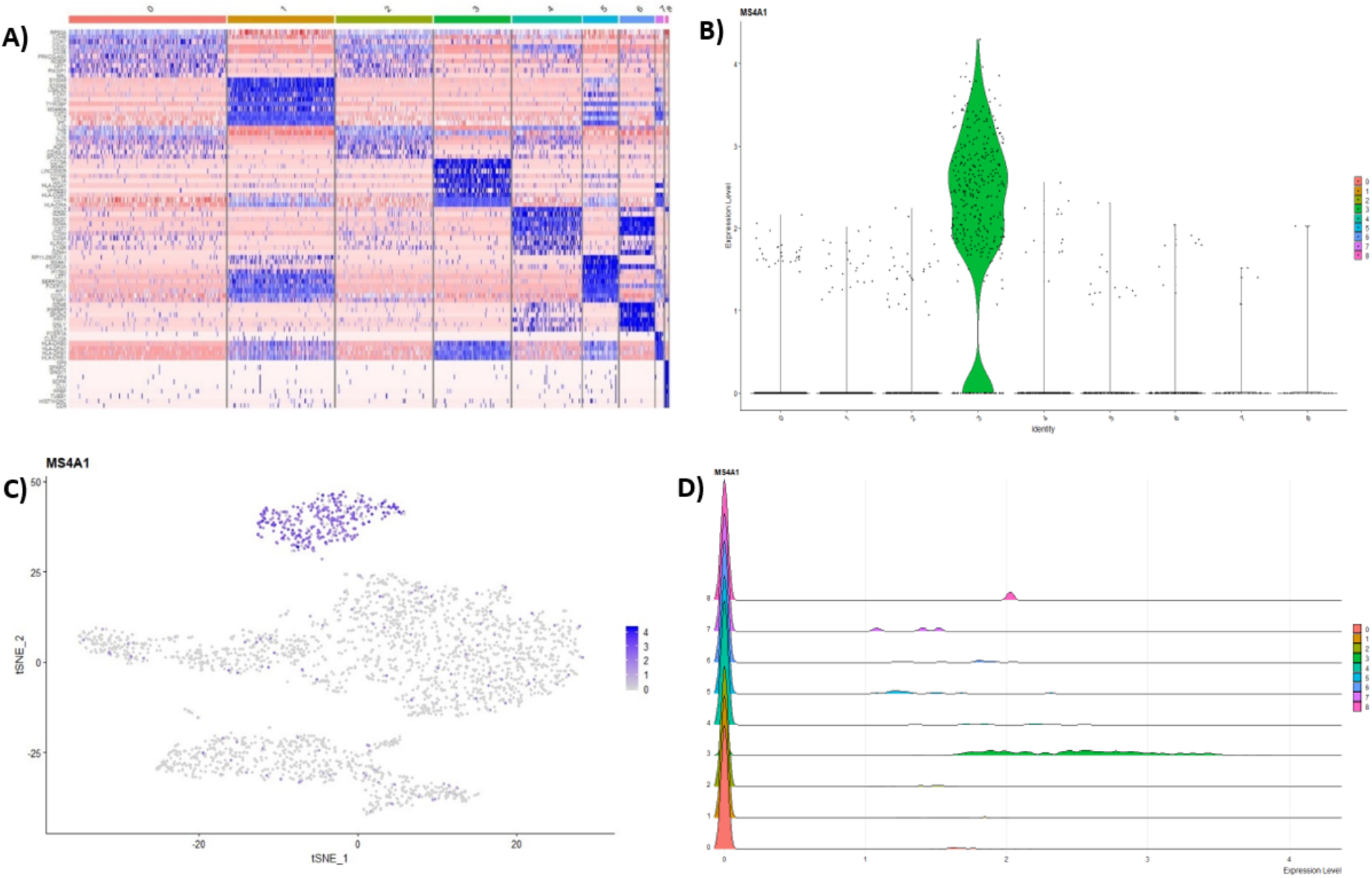
**A)** Heatmap showing the top 10 genes for each cluster in the 2700 PBMCs dataset, while Violin **B)**, Feature **C)**, and Ridge **D)** plots are shown for MS4A1 gene – a biomarker of B lymphocytes. Note that these DE visualization options are available in BingleSeq and are generated using Seurat’s inbuilt plotting functionality.

### Functional Annotation

Following DE analysis for both Bulk and scRNA-Seq pipelines, BingleSeq enables the extensive functional annotation analysis of DEGs using the GOseq pipeline (26). This is done in the ‘Functional Annotation’ tab where users can obtain results from KEGG pathway analysis and three types of GO categories, including ‘Cellular Component’, ‘Molecular Function’, and ‘Biological Function’ (**S5 Fig**). BingleSeq can also generate top 10 GO term histograms as well as to obtain additional information about a given GO term using the ‘GO.db’ package (27) (**S5 Fig**). Note that BingleSeq supports both Mouse and Human genomes (40,41).

### DE Package Comparison and Rank-Based Consensus

BingleSeq supplies an option to assess the agreement between the different DE analysis packages in the form of a Venn diagram. In the context of the Bulk RNA-Seq pipeline the overlap in DEGs is assessed on the results obtained using DESeq2, edgeR, and limma packages. For the case of scRNA-Seq this is done on three of Seurat’s inbuilt DE methods - MAST, Wilcoxon Rank Sum Test, and Student’s T test – see **S6A Fig**. Furthermore, BingleSeq provides a Rank-based consensus, as it was previously suggested as a mean to provide better confidence for DE results (43–45) – see **S6B Fig**.

### Inbuilt bulk and single cell RNA-Seq example datasets

BingleSeq features inbuilt test data for both Bulk and scRNA-Seq. Bulk RNA-Seq data is represented by a 3-sample contrast between HSV-1 infected control and interferon B treatment (38). The single-cell RNA-Seq example is a 10x Genomics public dataset looking at filtered data of 931 cells from a combined cortex, hippocampus and sub ventricular zone of an E18 mouse (https://support.10xgenomics.com/single-cell-gene-expression/datasets/2.1.0/neurons_900).

## Conclusions and Future Directions

BingleSeq is a comprehensive and intuitive solution that enables users to choose from multiple state-of-the-art Differential Expression analysis and unsupervised clustering packages according to their preferences or the dataset in question. In terms of Bulk RNA-Seq analyses, BingleSeq implements functionality that puts it close to, what is to our understanding, the best available similar applications – DEBrowser (30) and Omics Playground (31). In terms of scRNA-Seq, BingleSeq could be argued to be among the most exhaustive applications, such as ASAP (33) and singleCellTK (34). Thus, to our knowledge, BingleSeq is the only application to provide a solution for the analysis of both Bulk and scRNA-Seq data with multiple inbuilt state-of-the-art packages, alongside DEGs rank-based consensus and extensive functional annotation analysis.

Future work will focus on including more package options as well as extending and improving user control over the packages implemented in BingleSeq. Moreover, the implementation of both Bulk RNA-Seq and scRNA-Seq pipelines puts BingleSeq in a particularly good position to be the first package to implement user-friendly deconvolution of Bulk RNA-Seq results using scRNA-Seq data. Hence, an excellent and practical conclusion to the development of BingleSeq would be to include state-of-the-art deconvolution methods such as Cell Population Mapping (CPM) (46) or MUlti-Subject SIngle Cell deconvolution (MuSiC) (47).

BingleSeq is as an easy-to-install R package available as an archived zip file on GitHub at https://github.com/dbdimitrov/BingleSeq/. The application’s GitHub page provides an easy guide on how to install the application as well as examples of its general applicability and an extensive description of typical workflows when working with Bulk RNA-Seq and scRNA-Seq data.

## Acknowledgments

We acknowledge the financial support for bioinformatics developments as part of MRC (MC_UU_12014/12). We also extend our gratitude to Sejal Modha, Joseph Hughes, Richard Orton and Srikeerthana Kuchi for reviewing the manuscript prior to submission.

## Author Contributions

D.D. coded the application and contextualized the design of the pipelines. Q.G. conceptualized the idea for the application and supervised the project. Both D.D. and Q.G. contributed to the final version of the manuscript.

