## Supplementary material for "BingleSeq: A user-friendly R package for Bulk and Single-cell RNA-Seq Data Analysis": S1 File

### Bulk RNA-Seq Pipeline Evaluation

**Simulated data set evaluation**

Evaluation of BingleSeq's Bulk RNA-Seq pipeline was first performed using synthetic data generated with compcodeR package. compcodeR was used to simulate 6 two-condition datasets each with 20,000 features and a sequencing depth of 1e+07, varying between 0.7x and 1.4x this depth. There were no simulated outliers and of these 20,000 genes, 1000 were upregulated and 1000 downregulated. The datasets were divided in two groups that differed in the minimal differential expression strength of true positive DEGs. DEGs from the first and second groups had a minimal differential expression strength of 2 and 4, respectively. Each of these groups was composed of 3 datasets in total with 3, 6, and 12 replicates per condition.

BingleSeq was then used to generate DE results for each of the simulated datasets using the DE packages incorporated within its Bulk RNA-Seq pipeline. The obtained results were then assessed, and the sensitivity and specificity of the packages are shown in **fig. 1**. The lowest sensitivity was observed in the dataset with 3 replicates and effect size of 2, while the highest sensitivity was seen in the dataset with 12 replicates and DEGs with an effect size of 4 (**fig. 1A**). Therefore, as anticipated, sensitivity increased with both the number of replicates and effect size. Furthermore, the packages were rather conservative and relatively few DEGs were false positives, regardless of the dataset; hence, each package had high specificity (**fig. 1B**). These results confirm that the DE pipelines implemented in BingleSeq are capable of producing adequate DE analysis results.

Note that different sequencing depths were accounted for using the package-specific default methods for DESeq2 and edgeR, and voom and TMM normalization for limma. Similarly, the default test methods for each package were used for the analysis of the synthetic data set.

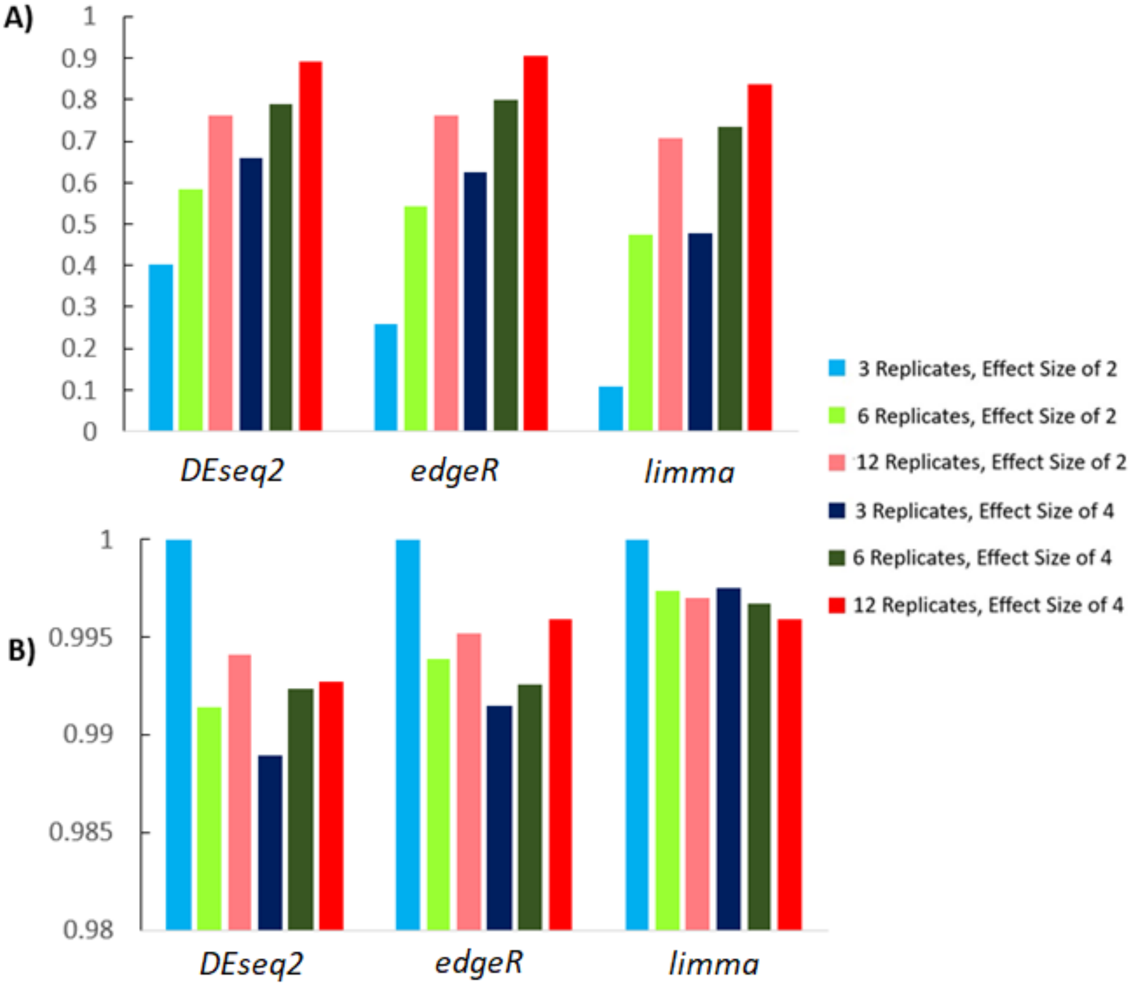


***Figure 1.*** *Statistical measures of performance (Sensitivity* ***A)*** *and Specificity* ***B)****) of DESeq2, edgeR, and limma as observed for each of the simulated datasets. See* ***Table 1*** *for tables with the obtained results.*

***Table 1**. Differential Analysis results for the simulated datasets obtained by **A)** DESeq2, **B)** edgeR, and **C)** limma pipelines as implemented in BingleSeq. Note that R stands for replicates and ES for effect size.*

***A)*** *DE analysis results obtained using DESeq2.*

| | 3R; ES2 | 6R; ES2 | 12R; ES2 | 3R; ES4 | 6R; ES4 | 12R; ES6 |
|---|---|---|---|---|---|---|
| DEGs | 894 | 1319 | 1629 | 1517 | 1718 | 1911 |
| True Positives | 744 | 1164 | 1522 | 1318 | 1580 | 1780 |
| False Positives | 0 | 155 | 107 | 199 | 138 | 131 |
| False Negatives | 1106 | 836 | 478 | 682 | 420 | 220 |
| True Negatives | 18000 | 17845 | 17893 | 17801 | 17862 | 17869 |
| Sensitivity | 0.402 | 0.582 | 0.761 | 0.659 | 0.790 | 0.890 |
| Specificity | 1.000 | 0.991 | 0.994 | 0.989 | 0.992 | 0.993 |

***B)*** *DE analysis results obtained using edgeR.*

| | 3R; ES2 | 6R; ES2 | 12R; ES2 | 3R; ES4 | 6R; ES4 | 12R; ES6 |
|---|---|---|---|---|---|---|
| DEGs | 575 | 1192 | 1606 | 1403 | 1730 | 1885 |
| True Positives | 494 | 1082 | 1519 | 1249 | 1596 | 1811 |
| False Positives | 0 | 110 | 87 | 154 | 134 | 74 |
| False Negatives | 1425 | 918 | 481 | 751 | 404 | 189 |
| True Negatives | 18000 | 17890 | 17913 | 17846 | 17866 | 17926 |
| Sensitivity | 0.257 | 0.541 | 0.760 | 0.625 | 0.798 | 0.906 |
| Specificity | 1.000 | 0.994 | 0.995 | 0.991 | 0.993 | 0.996 |

***C)*** *DE analysis results obtained using limma.*

| | 3R; ES2 | 6R; ES2 | 12R; ES2 | 3R; ES4 | 6R; ES4 | 12R; ES6 |
|---|---|---|---|---|---|---|
| DEGs | 218 | 995 | 1467 | 1000 | 1525 | 1749 |
| True Positives | 213 | 947 | 1413 | 955 | 1465 | 1675 |
| False Positives | 0 | 48 | 54 | 45 | 60 | 74 |
| False Negatives | 1782 | 1053 | 587 | 1045 | 535 | 325 |
| True Negatives | 18000 | 17952 | 17946 | 17955 | 17940 | 17926 |
| Sensitivity | 0.107 | 0.474 | 0.707 | 0.478 | 0.733 | 0.838 |
| Specificity | 1.000 | 0.997 | 0.997 | 0.998 | 0.997 | 0.996 |

### Real world data set evaluation

Following the evaluation using the synthetic data set, we used real world data (38) to showcase the applicability of BingleSeq's Bulk RNA-Seq pipeline. This dataset examined the differentially expressed genes (DEGs) between HSV-1 infected control and interferon B treatment.

First the data was filtered for genes with count per million (CPM) below 1 in at least 2 of the samples **(Fig. 2).**

**A)**

Raw Data Summary

| | Counts |
|---|---|
| mock_infected1_S1 | 31982285 |
| mock_infected2_S5 | 34742775 |
| mock_infected9_S9 | 28853256 |
| IFNb_treated12_S12 | 34376227 |
| IFNb_treated2_S8 | 40171321 |
| IFNb_treated4_S4 | 32451841 |
| Total Counts | 202577705 |
| Sample Median | 34376227 |
| Gene# | 58051 |

Showing 1 to 9 of 9 entries

Filtered Data Summary

| | Counts |
|---|---|
| mock_infected1_S1 | 31923291 |
| mock_infected2_S5 | 34682503 |
| mock_infected9_S9 | 28804742 |
| IFNb_treated12_S12 | 34321006 |
| IFNb_treated2_S8 | 40107936 |
| IFNb_treated4_S4 | 32402179 |
| Total Counts | 202241657 |
| Sample Median | 34321006 |
| Gene# | 13718 |

Showing 1 to 9 of 9 entries

**B)**

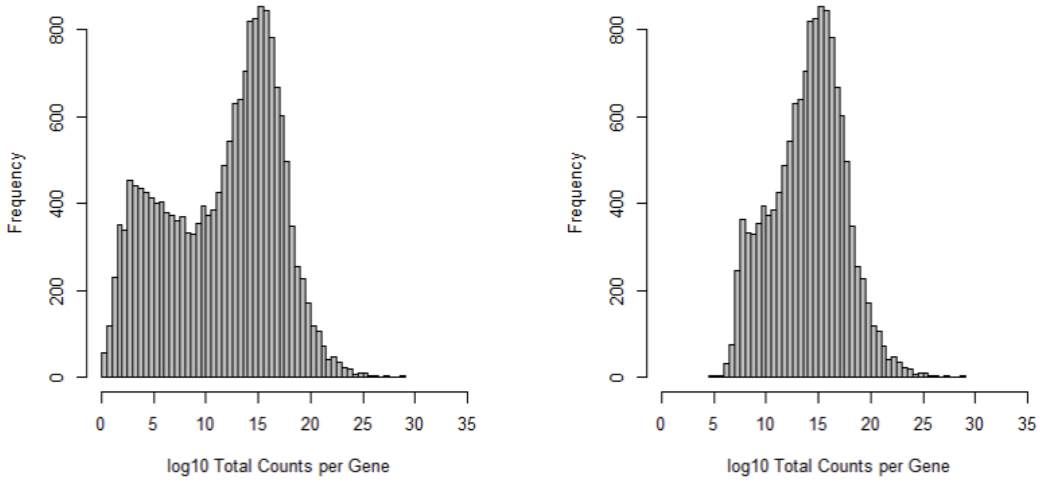


***Figure 2**. **A)** Count Data summaries before and after filtering with their corresponding **B)** gene frequency histograms.*

Following filtering differential expression (DE) testing was performed using limma with voom and TMM normalization. These results were then used for the subsequent visualization step of BingleSeq's Bulk RNA-Seq pipeline. In total, there were and 1053 up- and 1098 downregulated genes (**Fig. 3A**) with a visible distinction in the variation between the two conditions (**Fig. 3B**). Volcano (**Fig. 3C**) and MA (**Fig. 3D**) plots were then generated to represent the relationships between fold change (FC) versus significance and average expression, respectively.

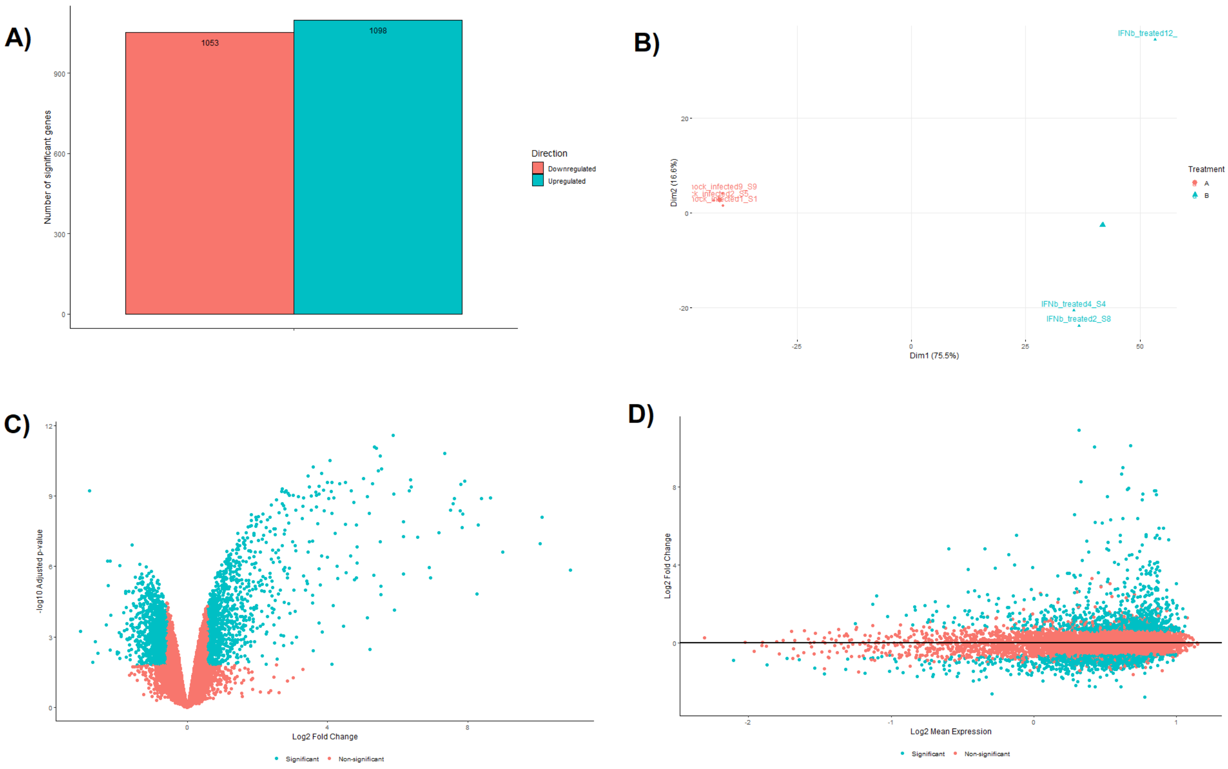


***Figure 3**. **A)** Barchart plot presenting the number of up- and downregulated genes **B)** PCA plot, **C)** Volcano plot, and **D)** MA plot.*

BingleSeq was also used to generate heatmaps for the top 20 up- and downregulated genes (**Fig. 4 A-B**) as well as for the functional gene annotation analysis of the DE results. The functional analysis revealed that 3 of the top 10 GO Biological Process terms were directly related to the cellular response to interferons. Thus, serving as further example of the applicability of the features implemented within our application. Note that each step was produced with DEGs filtered according to multiple-testing corrected p-value < 0.05 and log FC > 1.5.

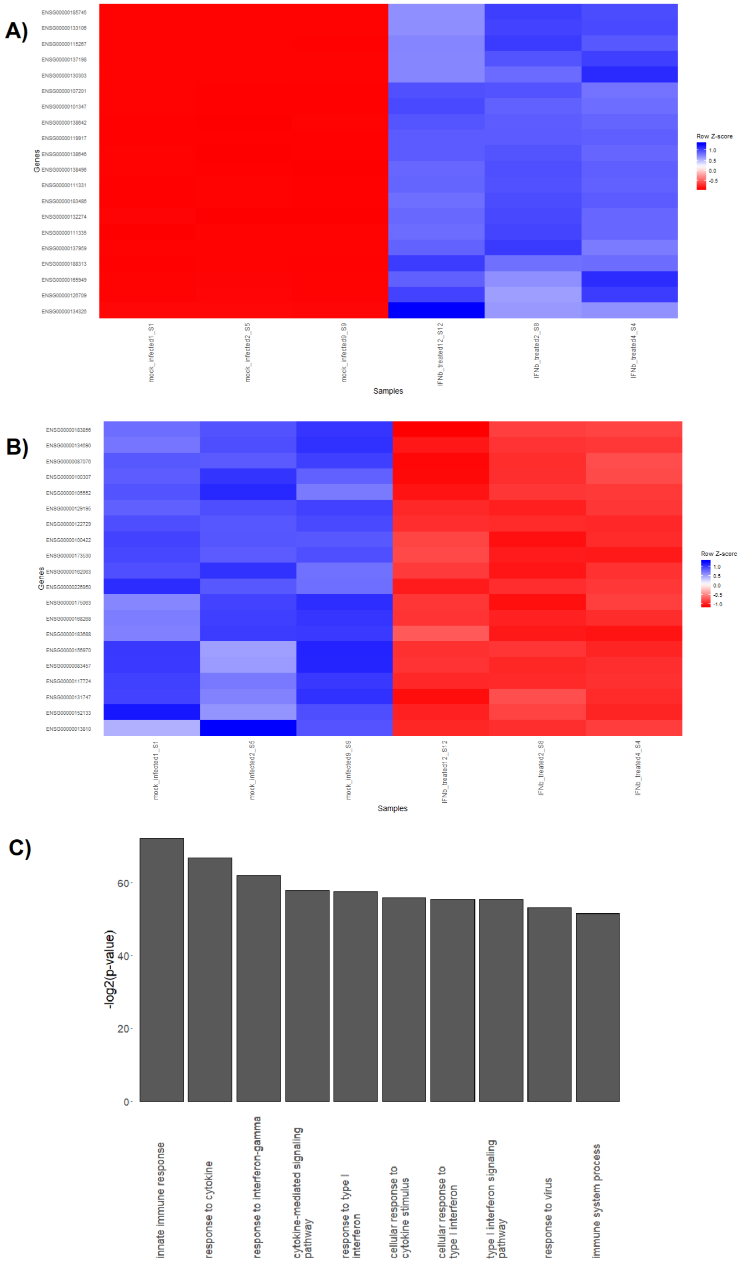


***Figure 4**. Heatmaps representing the top 20 **A)** up- and **B)** downregulated genes in the real-world data. **C)** Bar plot with showing the Top 10 most significant Biological Process Go terms.*

The agreement between DE packages was assessed by producing a Venn diagram with DEGs unfiltered by FC. The Venn diagram showed a considerable agreement between all 3 packages with DESeq2 and limma showing a slightly higher overlap in their results (**Fig. 5A**). Finally, the rank-based consensus table (**Fig. 5B**) showed considerable agreement in the ranking of the top 10 most significant genes, and it could also be used as an interactive tool that could be used to provide further confidence in specific genes of interest.

A)

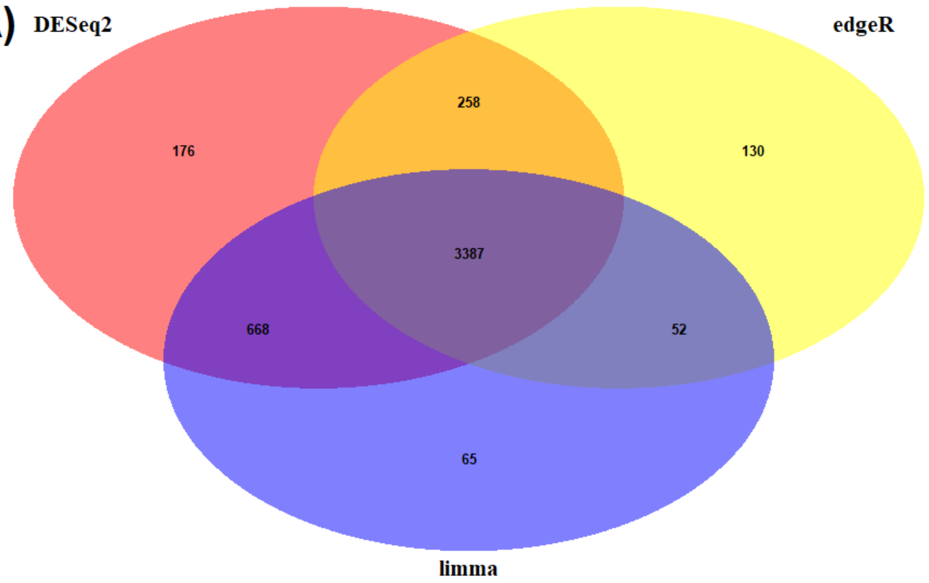


B)

Show 10 entries

Search:

| | edgeR Rank | edgeR adj.p-value | limma Rank | limma adj.p-value | DESeq2 Rank | DESeq2 adj.p-value | Ranking Consesus |
|---|---|---|---|---|---|---|---|
| ENSG00000183486 | 1 | 9.16556584892308e-301 | 4.5 | 5.03881799032289e-8 | 9 | 0 | 1 |
| ENSG00000119917 | 5 | 5.3094789581713e-250 | 1 | 3.32498473069603e-8 | 9 | 0 | 2 |
| ENSG00000107201 | 9 | 2.54644982000429e-219 | 2.5 | 4.05945330173205e-8 | 9 | 0 | 3 |
| ENSG00000111331 | 12 | 2.55002661384474e-214 | 2.5 | 4.05945330173205e-8 | 9 | 0 | 4 |
| ENSG00000111335 | 13 | 5.09825449235143e-213 | 4.5 | 5.03881799032289e-8 | 9 | 0 | 5 |
| ENSG00000137959 | 3 | 2.57292177332505e-288 | 16.5 | 2.06043462632089e-7 | 9 | 0 | 6 |
| ENSG00000138646 | 8 | 1.27523170607536e-228 | 16.5 | 2.06043462632089e-7 | 9 | 0 | 7 |
| ENSG00000115267 | 16 | 1.87859083393152e-209 | 9 | 1.1905802584263e-7 | 9 | 0 | 8 |
| ENSG00000138642 | 18 | 1.49283520021086e-205 | 8 | 1.14784524067696e-7 | 9 | 0 | 9 |
| ENSG00000133106 | 19 | 3.60431133203377e-189 | 12 | 1.9354154233462e-7 | 9 | 0 | 10 |

***Figure 5**. **A)** Venn diagram showing the overlap of DEGs obtained by the scRNA-Seq pipeline for the real-world data set. **B)** An interactive rank-based consensus table showing the top 10 highest ranked genes generated with the same data.*
