## Supplementary material for "BingleSeq: A user-friendly R package for Bulk and Single-cell RNA-Seq Data Analysis": S2 File

### scRNA-Seq Pipeline Evaluation

The evaluation of BingleSeq's scRNA-Seq pipeline was performed by reproducing and extending the results of Seurat's online tutorial (https://satijalab.org/seurat/v3.0/pbmc3k_tutorial.html) . The tutorial is based on a 10x Genomics dataset of 2700 Peripheral Blood Mononuclear Cells (PBMCs) with ~69,000 reads per cell. This tutorial makes use of a 10x Genomics dataset of 2700 Peripheral Blood Mononuclear Cells (PBMCs) with ~69,000 reads per cell (https://support.10xgenomics.com/single-cell-gene-expression/datasets/1.1.0/pbmc3k).
To evaluate BingleSeq's applicability and reproducibility, this evaluation followed strictly the Seurat's tutorial and the parameters used in it.
First, cells with unique gene counts less than 200 and above 2500 were filtered (**Fig. 1**).

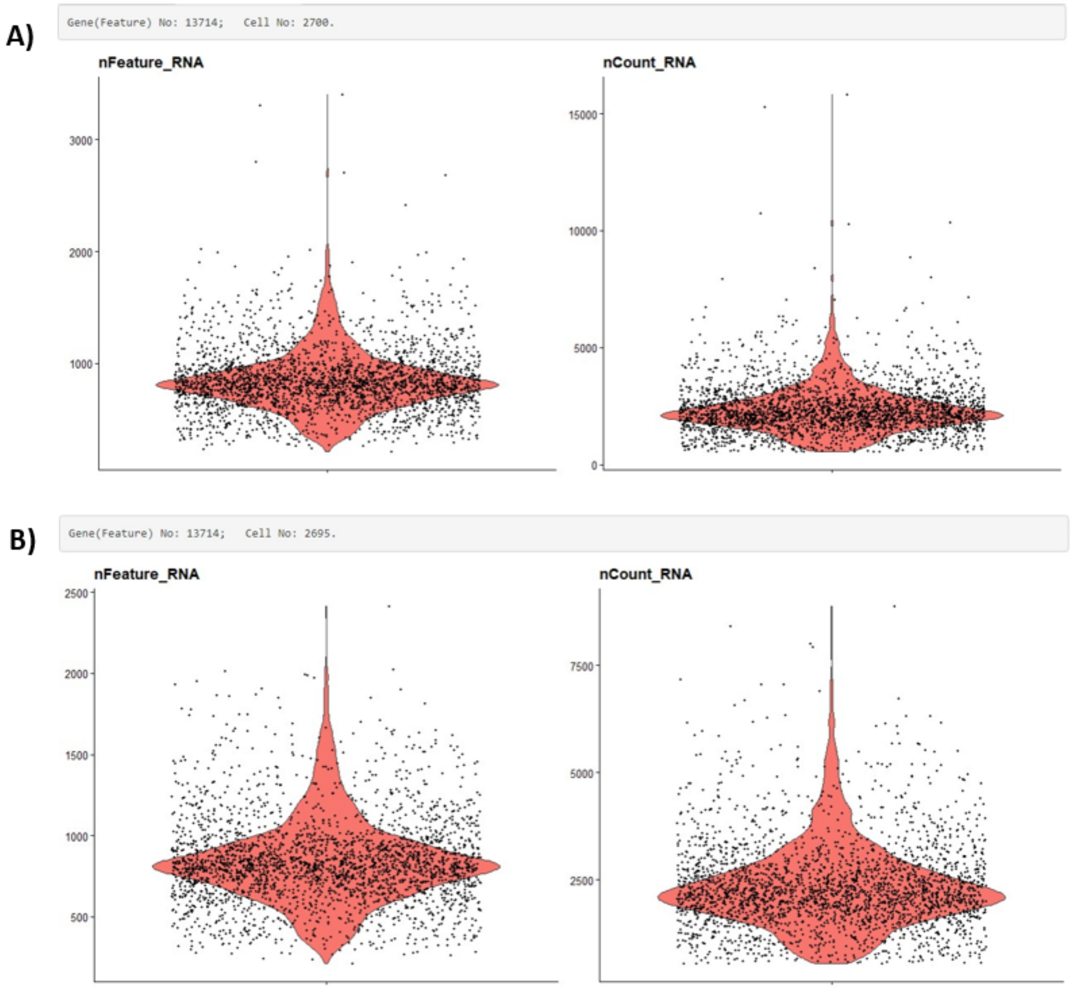


***Figure 1.*** *Violin plots and Feature/RNA count number summary produced as part of BingleSeq's cell outlier filtering procedure. The Violin plots are presented* ***A)*** *before and* ***B)*** *after outlier filtering of the PBMC dataset. Cells are filtered according to the number of expressed features per cell (nFeature), while nCount_RNA represents the number of UMIs.*

Following Quality Control, data normalization was performed using the "LogNormalize" method with a scale factor of 10,000. Subsequent to normalization, feature selection was performed using the "vst" procedure and the top 2000 most variable genes were selected for downstream analysis with Seurat's method (**Fig. 2**).

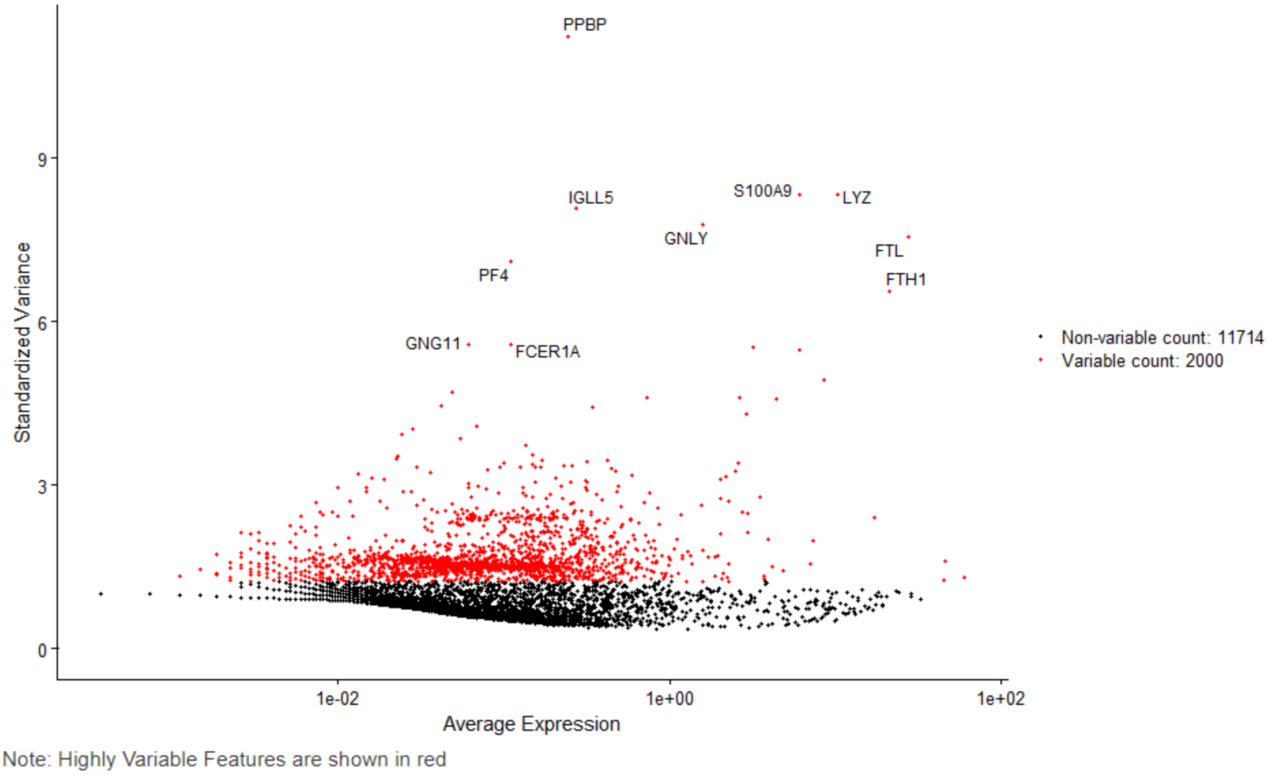


***Figure 2***. *Variable Features plot generated following normalization and feature selection. Note that the top 2000 most variable genes are coloured in red and the top 10 most variable genes are also labelled.*

Subsequently, the data was scaled, linear dimensional reduction was performed, and the true dimensionality of the dataset was determined using an elbow plot (**Fig. 3A**). In an analogous manner to the tutorial, the elbow was observed around the 9-10$^{th}$ PC. Hence, this was the dimensionality used in unsupervised clustering. See **Fig. 3B-C** for further exploration of the dimensionality of the dataset using PC heatmap.

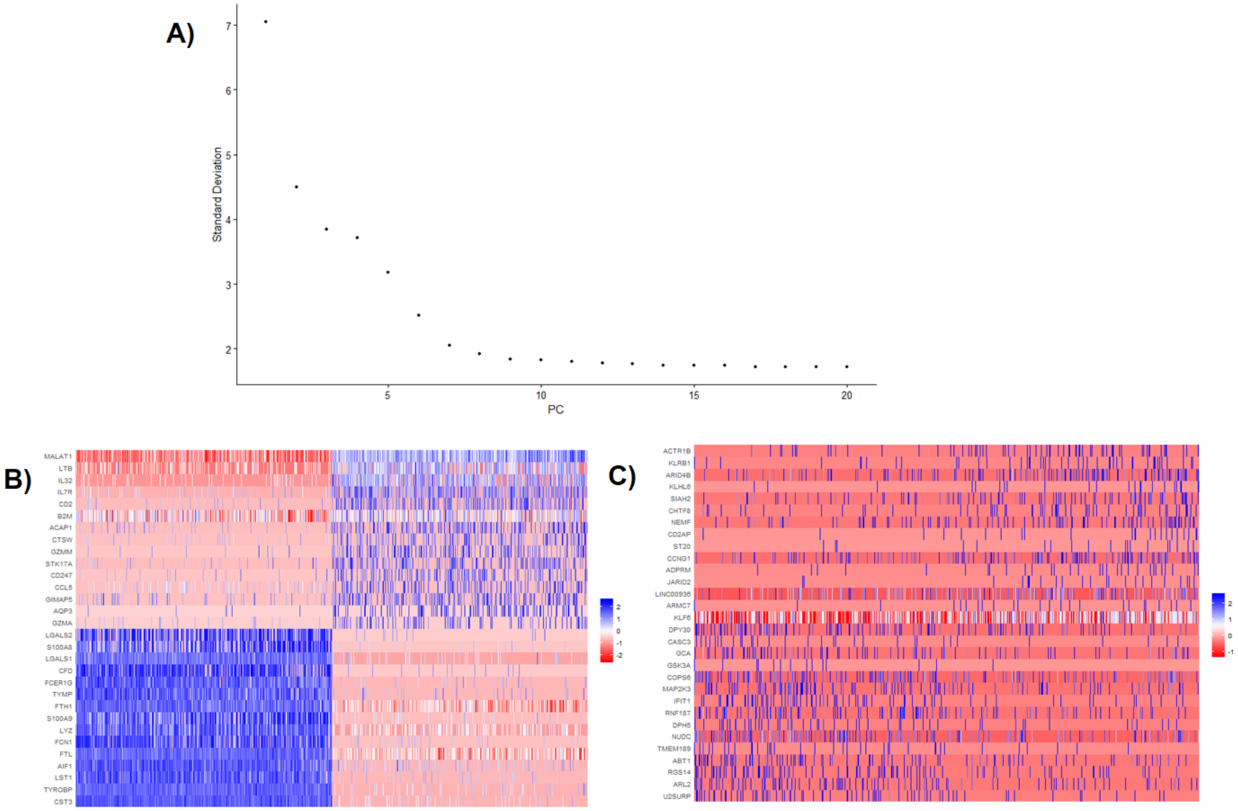


***Figure 3**. **A)** Elbow plot produced for the ~2700 PBMCs dataset and subsequently used to determine the its true dimensionality. **B)** 1$^{st}$ PC Heatmap with the top 10 most variable Genes which is very likely to represent the true dimensionality of the dataset. In contrast, C) is the 15$^{th}$ PC Heatmap which is unlikely to represent true dimensionality.*

Unsupervised clustering was performed with Seurat, monocle, and SC3. Clustering with Seurat was performed with dimensionality and resolution parameters identical to those used in the tutorial and yielded analogous results (**Fig. 4A** and **4D**). Unsupervised clustering with monocle and SC3 was performed by explicitly setting the number of clusters to 9 which resulted in cells being clustered in a highly analogous way to Seurat's results (**Fig. 4B-C**).

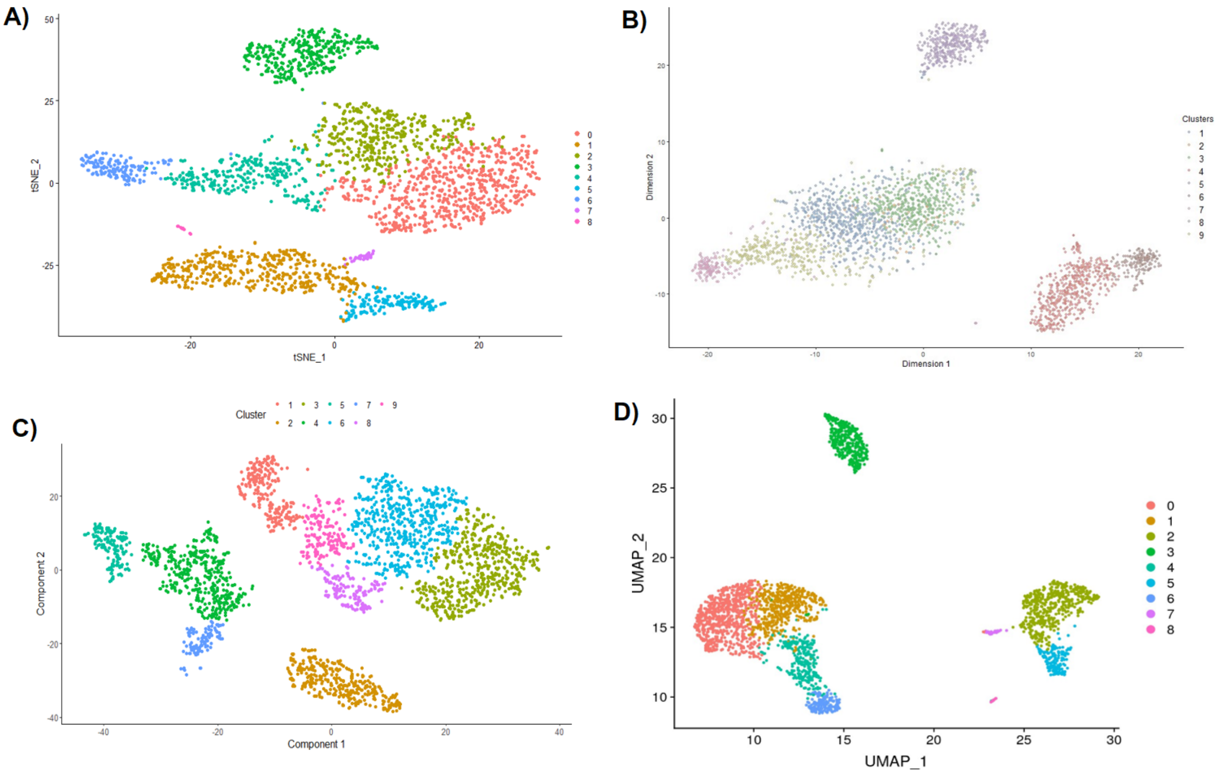


***Figure 4***. *tSNE plots generated for the 2700 PBMCs dataset using* ***A)*** *Seurat,* ***B)*** *SC3, and* ***C)*** *monocle.* ***D)*** *Unsupervised clustering results obtained by Satija lab using UMAP dimensionality reduction for the same dataset.*

Furthermore, a similar analysis was conducted using a larger 10x Genomics dataset composed of ~5400 PBMCs with a mean sequencing depth of ~28,000 reads per cell (https://support.10xgenomics.com/single-cell-gene-xpression/datasets/1.1.0/pbmc6k).

The obtained clustering results (**Fig. 5**) were highly similar to the ones observed in the smaller PBMCs dataset. Thus, further confirming the applicability of BingleSeq's scRNA-Seq pipeline.

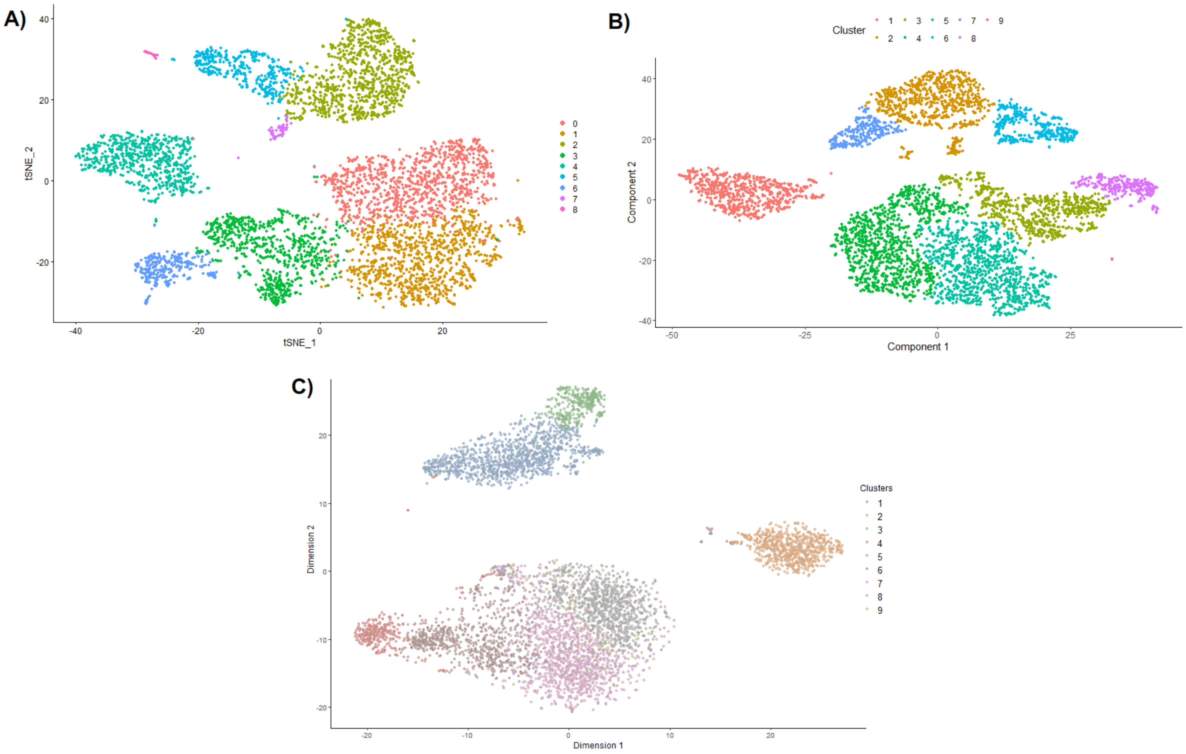


***Figure 5**. **A)** Seurat **B)** SC3, **C)** monocle unsupervised clustering results for the 10x Genomics dataset composed of ~5400 PBMCs. Note that cluster number was explicitly set to 9 for SC3 and monocle.*

Subsequent to clustering, DE analysis of all clusters was performed in an analogous manner to the tutorial using the Wilcoxon rank sums test. In turn, this yielded matching results (**Fig. 6A**).

MS4A1 is a B lymphocyte marker gene and was hence chosen to pinpoint the cluster corresponding to B cells (**Fig. 6B-D**); thus, confirming BingleSeq's applicability to yield meaningful DE results and their subsequent use in identifying cluster identity.

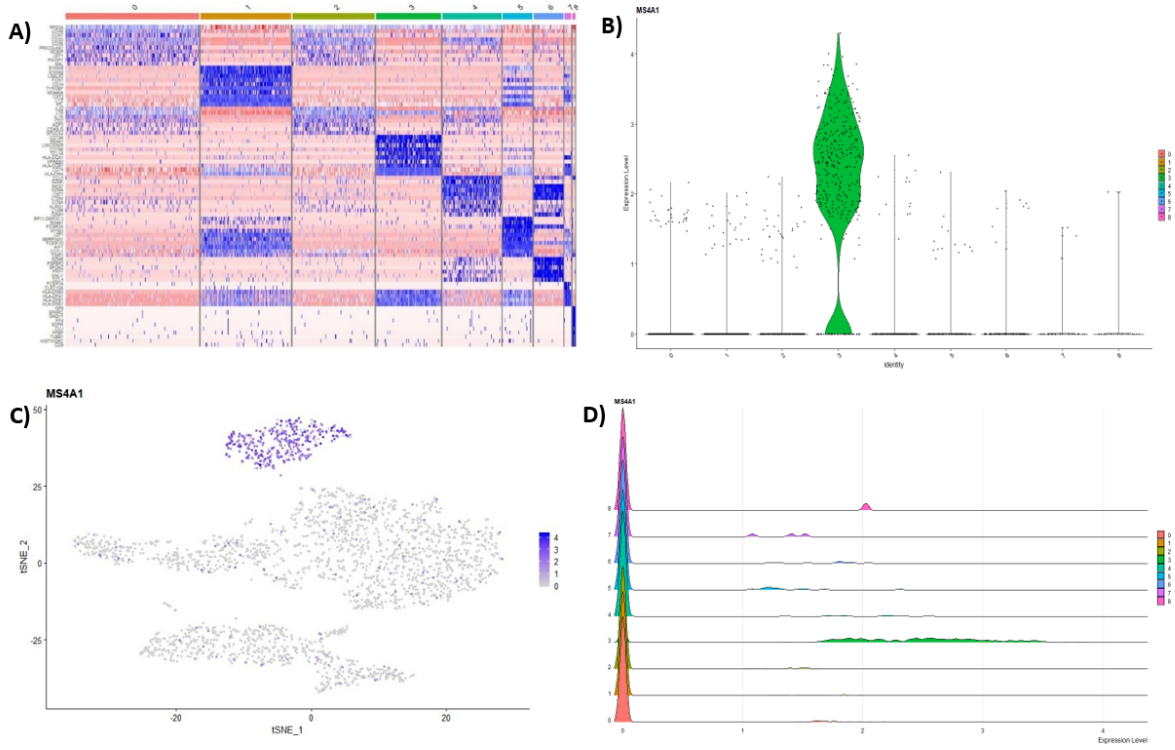


**Fig 6. A)** Heatmap showing the top 10 genes for each cluster in the 2700 PBMCs dataset, while Violin **B)**, Feature **C)**, and Ridge **D)** plots are shown for MS4A1 gene – a biomarker of B lymphocytes. Note that these DE visualization options are available in BingleSeq and are generated using Seurat's inbuilt plotting functionality.

Following DE analysis, BingleSeq's 'Functional Annotation' tab was used to gain further insight about the clusters. The functional annotation analysis further confirmed that cluster 3 corresponds to B lymphocytes, as its most significant GO Term as well as 3 of its top 20 Biological processes were specifically associated with B lymphocytes (**Fig. 7**). Thus, serving as proof for the applicability of BingleSeq's Functional Annotation pipeline in revealing crucial phenotypic insight.

A)

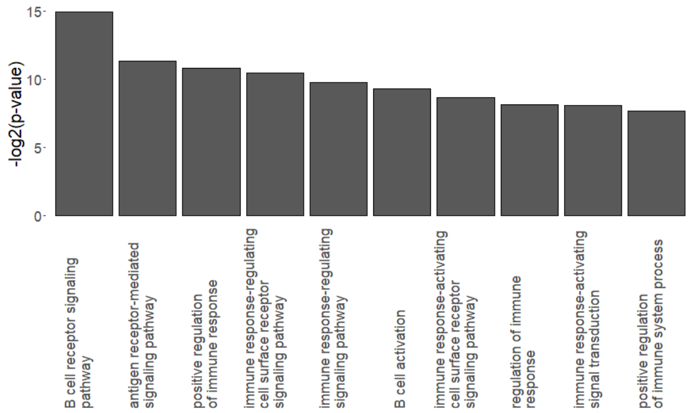


B)

DE Genes Table | GO Term Table | GO Term Histogram | GO Term Info

Show 25 entries

Search:

| | category | over_represented_pvalue | under_represented_pvalue | numDEInCat | numInCat | term | ontology |
|---|---|---|---|---|---|---|---|
| 8514 | GO:0050853 | 0.00003194 | 1.000 | 6 | 50 | B cell receptor signaling pathway | BP |
| 8512 | GO:0050851 | 0.0003801 | 1.000 | 8 | 208 | antigen receptor-mediated signaling pathway | BP |
| 8475 | GO:0050778 | 0.0005444 | 1.000 | 12 | 704 | positive regulation of immune response | BP |
| 861 | GO:0002768 | 0.0006975 | 1.000 | 9 | 353 | immune response-regulating cell surface receptor signaling pathway | BP |
| 858 | GO:0002764 | 0.001148 | 1.000 | 10 | 510 | immune response-regulating signaling pathway | BP |
| 6359 | GO:0042113 | 0.001600 | 1.000 | 7 | 207 | B cell activation | BP |
| 645 | GO:0002429 | 0.002474 | 1.000 | 8 | 320 | immune response-activating cell surface receptor signaling pathway | BP |
| 8473 | GO:0050776 | 0.003562 | 1.000 | 12 | 914 | regulation of immune response | BP |
| 851 | GO:0002757 | 0.003611 | 1.000 | 9 | 476 | immune response-activating signal transduction | BP |
| 793 | GO:0002684 | 0.004775 | 1.000 | 12 | 967 | positive regulation of immune system process | BP |
| 791 | GO:0002682 | 0.008316 | 1.000 | 14 | 1424 | regulation of immune system process | BP |
| 1828 | GO:0006955 | 0.008316 | 1.000 | 16 | 1854 | immune response | BP |
| 535 | GO:0002253 | 0.008599 | 1.000 | 9 | 551 | activation of immune response | BP |
| 8216 | GO:0048584 | 0.02433 | 1.000 | 16 | 2102 | positive regulation of response to stimulus | BP |
| 532 | GO:0002250 | 0.03000 | 1.000 | 7 | 355 | adaptive immune response | BP |
| 6352 | GO:0042100 | 0.03000 | 1.000 | 4 | 71 | B cell proliferation | BP |
| 617 | GO:0002376 | 0.04437 | 1.000 | 18 | 2727 | immune system process | BP |

***Figure 7**. **A)** Histogram of the Top 10 Biological Process Go Terms for cluster 3 and **B)** a table for the Top 20 Biological Process Go Terms.*

Finally, BingleSeq's 'DE Comparison' tab was used to assess the agreement between the different differential gene expression (DE) analysis methods implemented within Seurat's pipeline. The results showed a high-level of overlap and agreement between the different DE methods (**Fig 9A**), with Wilcoxon and MAST having a particularly high agreement. Furthermore, the interactive Rank-based consensus table can be used to obtain further confidence for the significance of specific features of interest (**Fig 8B**).

A)

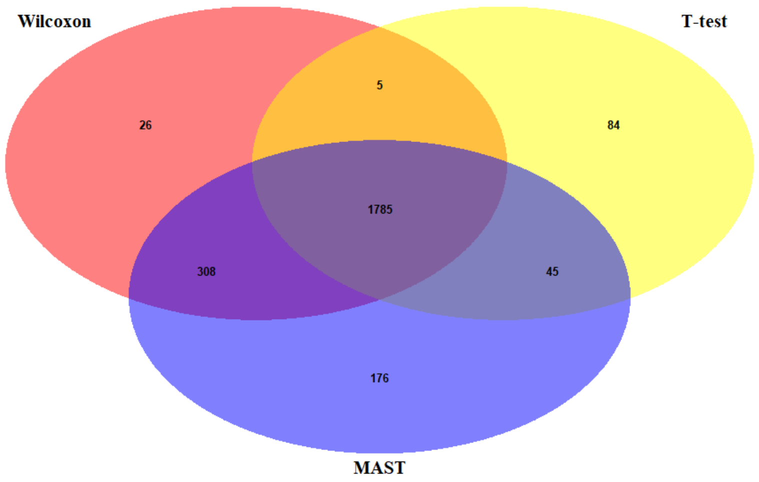


B)

Show 10 entries

Search:

| | Wilcoxon Rank | Wilcoxon adj.p-value | T-test Rank | T-test adj.p-value | MAST Rank | MAST adj.p-value | Ranking Consesus |
|---|---|---|---|---|---|---|---|
| CFP.1 | 97 | 2.5301454771267e-152 | 32 | 1.22694339616222e-222 | 82 | 4.38911743767198e-130 | 1 |
| FCN1.1 | 115 | 5.88190462390935e-142 | 236 | 1.91890212280612e-83 | 6 | 0 | 2 |
| TYMP | 159 | 4.96774269310204e-122 | 69 | 1.21220978903259e-178 | 245 | 6.77514268905079e-76 | 3 |
| MS4A6A | 74 | 4.35437699471525e-169 | 427 | 1.97184168391075e-47 | 29 | 2.07482617382848e-219 | 4 |
| CSF1R | 61 | 4.5143279324225e-185 | 130 | 1.72856293719667e-116 | 376 | 1.11259516710577e-56 | 5 |
| RPL23A | 240 | 1.13201215939462e-95 | 302 | 1.02748072312992e-67 | 45 | 1.99689110922324e-176 | 6 |
| RPS15A | 254 | 1.0947034972274e-92 | 206 | 7.76662674357972e-91 | 145 | 2.35017022301052e-99 | 7 |
| CD74.2 | 295 | 1.8192992694437e-83 | 307 | 8.58008635919608e-67 | 6 | 0 | 8 |
| ALDH2 | 112 | 1.73857771420774e-142 | 373 | 6.06047671285392e-54 | 130 | 4.04024967713762e-107 | 9.5 |
| CPVL | 249 | 5.11325442579919e-94 | 132 | 2.02326222241065e-114 | 234 | 1.8552740816104e-77 | 9.5 |

Showing 1 to 10 of 2,314 entries

Previous 1 2 3 4 5 … 232 Next

Download Ranking Consensus

***Figure 8. A)*** *Venn diagram showing the overlap of DE results obtained using 3 selected methods and a* ***B)*** *an interactive rank-based consensus table.*
