## Supporting information for "BingleSeq: A user-friendly R package for Bulk and Single-cell RNA-Seq Data Analysis"

### Supplementary materials

**S1 Table**. Comparison between *BingleSeq*'s Bulk RNA-Seq pipeline and other similar applications.

| **Functionality** | ***BingleSeq*** | *DEapp* | *DEBrowser* | *Omics Playground* |
|---|---|---|---|---|
| Filter Low Gene Counts | ✓ | ✓ | ✓ | ✓ |
| Batch-effect Correction | ✓ | | ✓ | ✓ |
| Implements *DESeq2* | ✓ | ✓ | ✓ | ✓ |
| Implements *EdgeR* | ✓ | ✓ | ✓ | ✓ |
| Implements *limma* | ✓ | ✓ | ✓ | ✓ |
| Additional DE methods | | | | ✓ |
| PCA plot | ✓ | ✓ | ✓ | ✓ |
| Summary Barchart | ✓ | | | ✓ |
| MA plot | ✓ | | ✓ | ✓ |
| Volcano plot | ✓ | ✓ | ✓ | ✓ |
| All to all scatter plot | | | ✓ | ✓ |
| Heatmap | ✓ | | ✓ | ✓ |
| Interquartile Range plot | | ✓ | ✓ | ✓ |

**S2 Table**. Comparison between *BingleSeq*'s scRNA-Seq pipeline and other similar applications.

| **Functionality** | ***BingleSeq*** | *singleCellTK* | *SeuratWizard* | *ASAP* | *Omics Playground* |
|---|---|---|---|---|---|
| Filtering/Normalization/Scaling | ✓ | ✓ | ✓ | ✓ | ✓ |
| Batch effect correction | | ✓ | | | ✓ |
| Pre-cluster Noise Filtering | ✓ | | ✓ | ✓ | |
| PCA Dimension Reduction | ✓ | ✓ | ✓ | ✓ | ✓ |
| tSNE Dimension Reduction | ✓ | ✓ | ✓ | ✓ | ✓ |
| UMAP Dimension Reduction | | ✓ | | | |
| Multiple Clustering Algorithms | ✓ | ✓ | ✓ | ✓ | |
| Estimate cluster number | ✓ | ✓ | | ✓ | ✓ |
| Define Cluster Number | ✓ | ✓ | | ✓ | |
| Trajectory (Pseudotime) | | ✓ | | ✓ | |
| DGE/Biomarker Visualization | ✓ | ✓ | ✓ | | ✓ |
| Differential Expression | ✓ | ✓ | ✓ | ✓ | ✓ |

**S3 Table**. Comparison between *BingleSeq*'s additional features and other similar applications.

| **Functionality** | ***BingleSeq*** | *singleCell TK* | *DEapp* | *DEBrowser* | *SeuratWizard* | *ASAP* | *Omics Playground* |
|---|---|---|---|---|---|---|---|
| Customizable plots | ✓ | | | ✓ | ✓ | ✓ | ✓ |
| Interactive plots | | | | ✓ | ✓ | ✓ | ✓ |
| GO Term Analysis | ✓ | | | ✓ | ✓ | ✓ | ✓ |
| GO Term Search | ✓ | | | | | | |
| KEGG Pathway Analysis | ✓ | | | ✓ | ✓ | ✓ | ✓ |
| Disease-related Analysis | | | | ✓ | ✓ | | |
| Oncogenic Signatures | | | | | | ✓ | |
| Copy number variation analysis | | | | | | | ✓ |
| Interactive Network Visualizations | | | | | | | ✓ |
| DE Method Comparison | ✓ | ✓ | ✓ | | | | ✓ |
| DE Rank-based Consensus | ✓ | | | | | | |
| Intersection Analysis | | | | | | | ✓ |
| Complex Experimental Design | ✓ | ✓ | ✓ | ✓ | ✓ | | ✓ |
| Download data and plots | ✓ | | | ✓ | ✓ | ✓ | ✓ |
| Inbuilt scRNA-Seq Pipeline | ✓ | | | | | ✓ | ✓ |
| Proteomics Analysis | | | | | | | ✓ |
| Manual cluster annotation | | | | | | ✓ | |
| Development | R/shiny | R/shiny | R/shiny | R/shiny | R/shiny | Multi-language | R/shiny |

A)

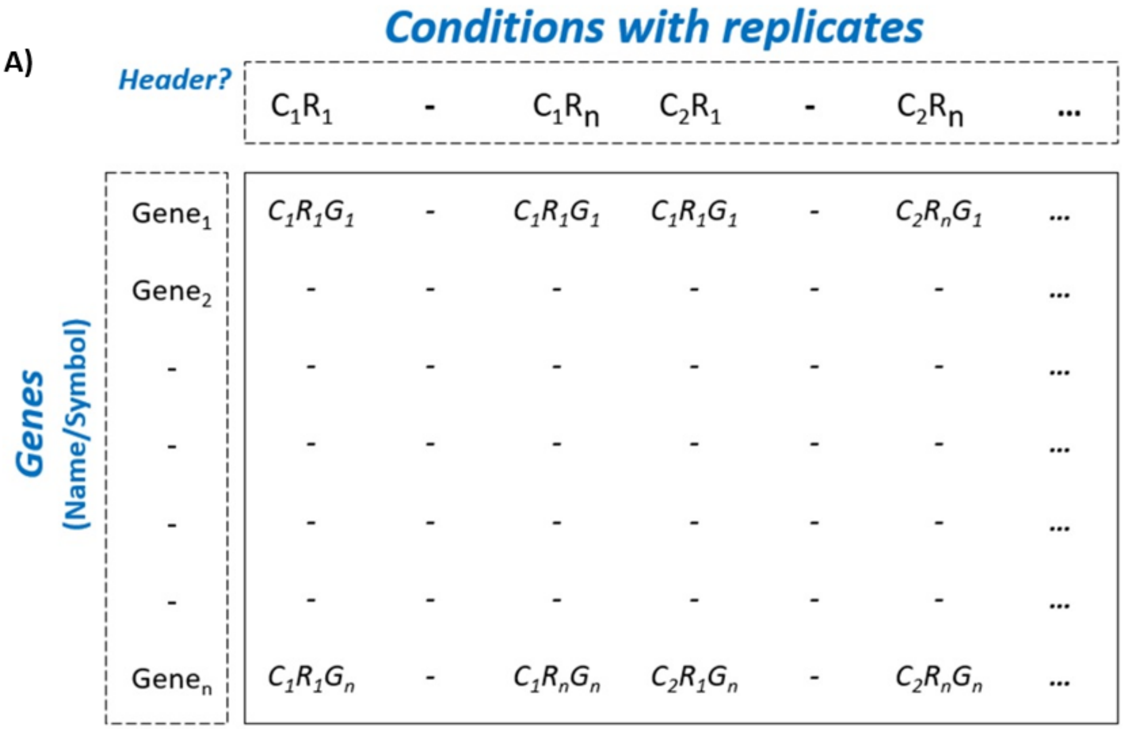


B)

| sample | batch | treatment |
|---|---|---|
| wt_rep1 | 1 | A |
| wt_rep2 | 1 | A |
| wt_rep3 | 2 | A |
| wt_rep4 | 2 | A |
| mutant_rep1 | 1 | B |
| mutant_rep2 | 1 | B |
| mutant_rep3 | 2 | B |
| mutant_rep4 | 2 | B |

***S1 Fig. A)*** *Gene count table and* ***B)*** *metadata table formats required by BingleSeq's Bulk RNA-Seq pipeline. The use of metadata tables was inspired by similar applications preceding BingleSeq - DEapp and DEBrowser.*

A)

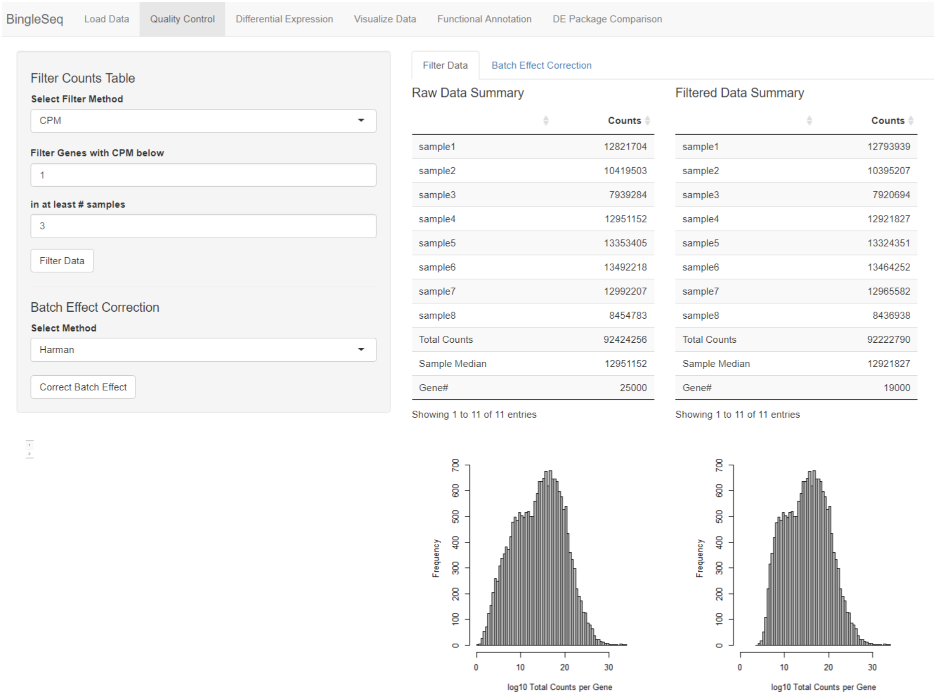


B)

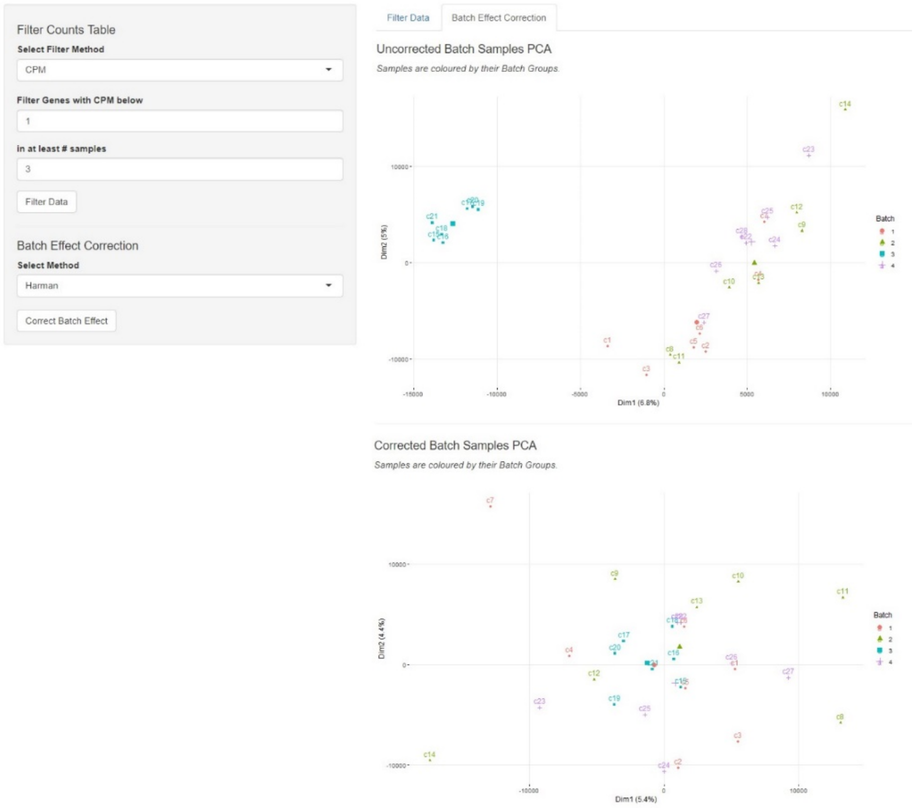


***S2 Fig.*** *An overview of BingleSeq's user interface (UI) and the 'Quality Control' tab with* ***A)*** *'Filter Data' and* ***B)*** *'Batch Effect Correction' subtabs.*

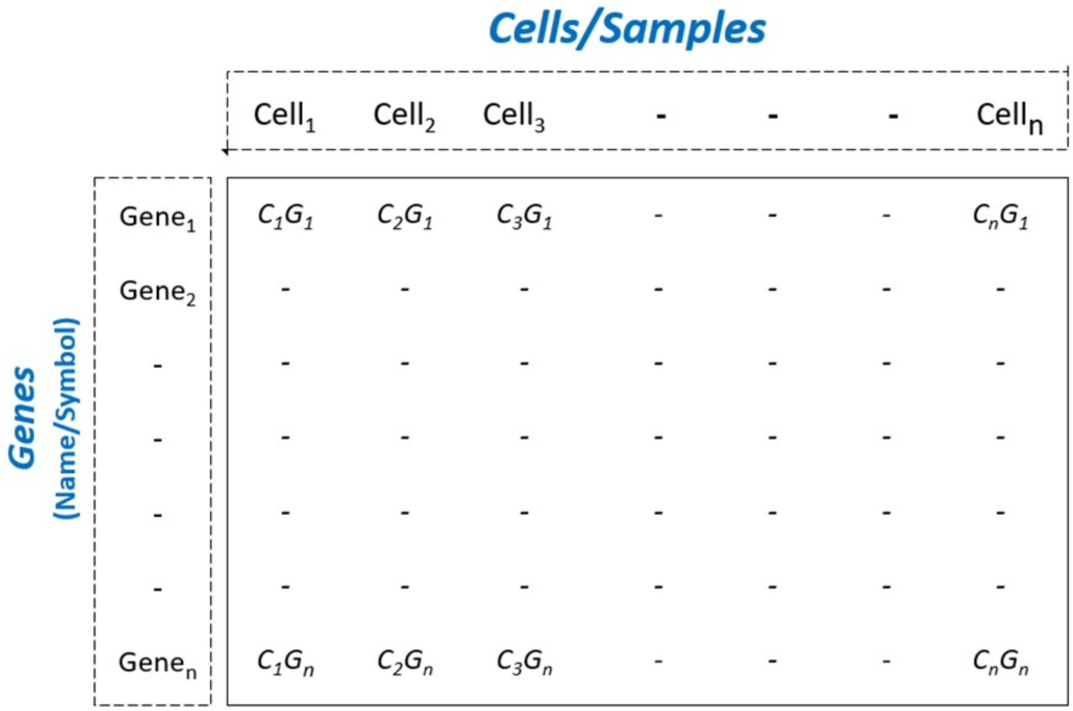


***S3 Fig.*** *Gene counts table format required by BingleSeq's scRNA-Seq pipeline.*

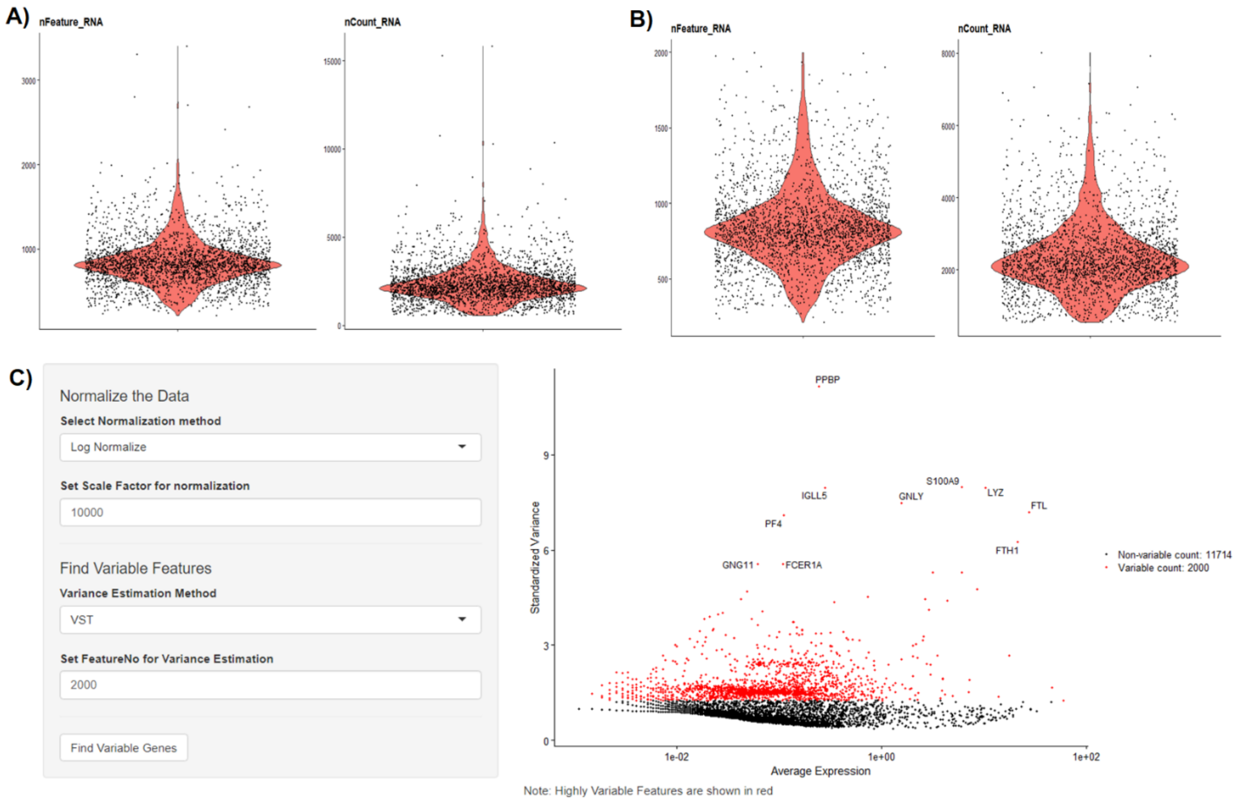


***S4 Fig.*** *Violin plots representing unfiltered (**A**) and filtered (**B**) scRNA-seq data and C) BingleSeq's Normalization tab with a variable features plot showing the top 2000 most variable genes.*

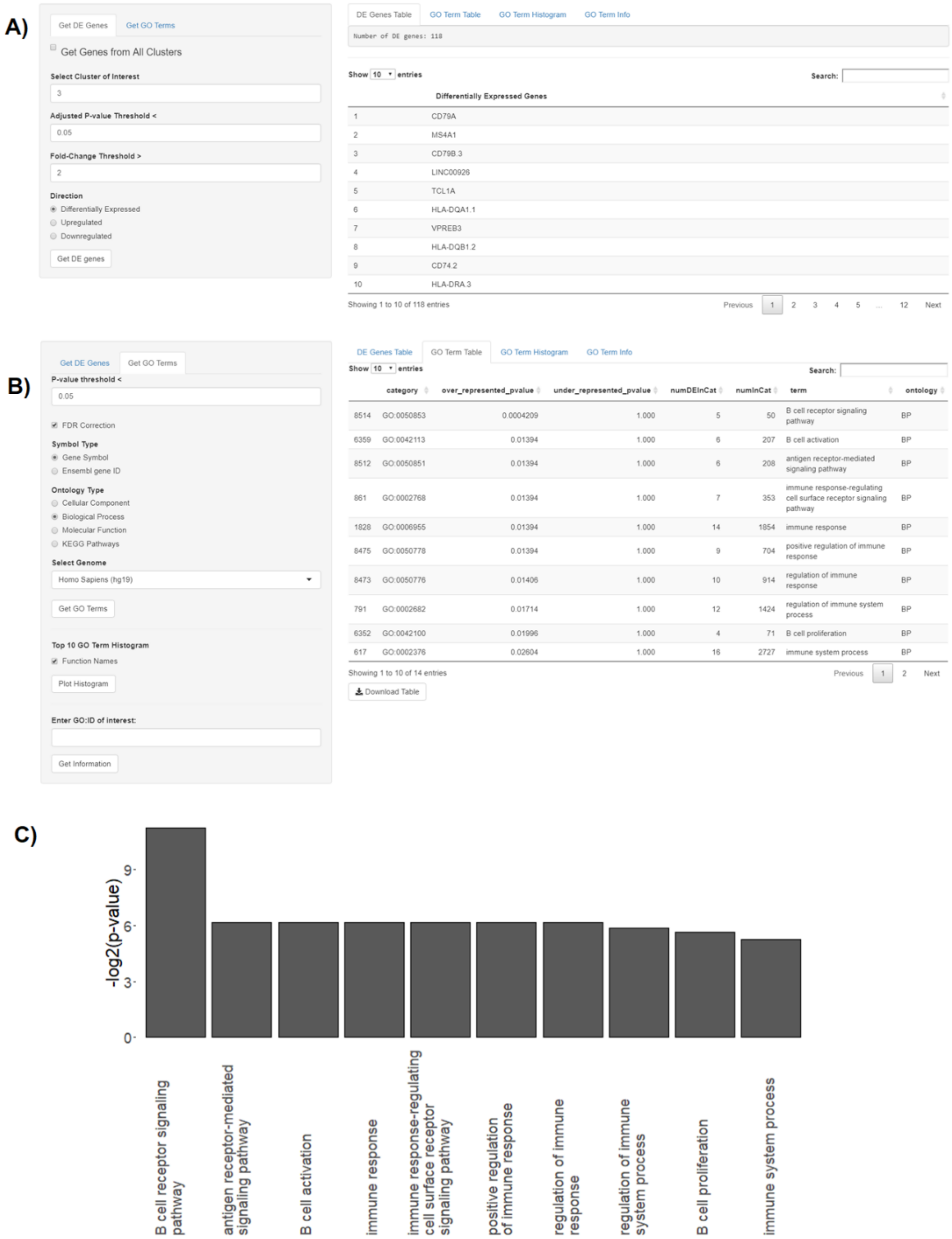


***S5 Fig***. *Overview of the 'Functional Annotation' tab with the* ***A)*** *'Get DE Genes' and* ***B)*** *'Get GO terms' control panels shown with their corresponding output. The latter shows the GO terms associated with Biological processes identified using DEGs from cluster 3 which as identified by the high expression of MS4A1 gene corresponds to the B cell phenotype.* ***C)*** *Top 10 GO terms histogram showing 3 of the top 10 processes belonging to B cell activation. The data DGEs presented here were generated with Seurat's pipeline in accordance with its tutorial - https://satijalab.org/seurat/v3.0/pbmc3k_tutorial.html.*

A)

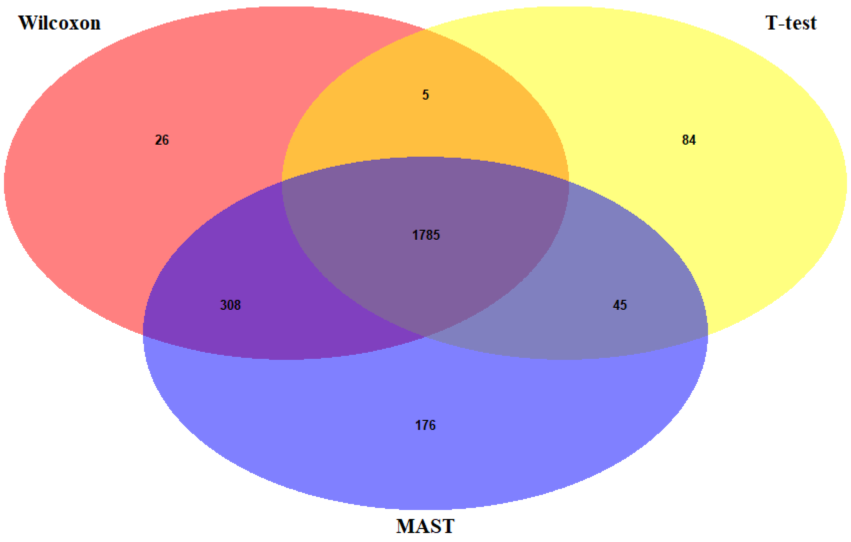


B)

Show 10 entries

Search:

| | Wilcoxon Rank | Wilcoxon adj.p-value | T-test Rank | T-test adj.p-value | MAST Rank | MAST adj.p-value | Ranking Consesus |
|---|---|---|---|---|---|---|---|
| CFP.1 | 97 | 2.5301454771267e-152 | 32 | 1.22694339616222e-222 | 82 | 4.38911743767198e-130 | 1 |
| FCN1.1 | 115 | 5.88190462390935e-142 | 236 | 1.91890212280612e-83 | 6 | 0 | 2 |
| TYMP | 159 | 4.96774269310204e-122 | 69 | 1.21220978903259e-178 | 245 | 6.77514268905079e-76 | 3 |
| MS4A6A | 74 | 4.35437699471525e-169 | 427 | 1.97184168391075e-47 | 29 | 2.07482617382848e-219 | 4 |
| CSF1R | 61 | 4.5143279324225e-185 | 130 | 1.72856293719667e-116 | 376 | 1.11259516710577e-56 | 5 |
| RPL23A | 240 | 1.13201215939462e-95 | 302 | 1.02748072312992e-67 | 45 | 1.99689110922324e-176 | 6 |
| RPS15A | 254 | 1.0947034972274e-92 | 206 | 7.76662674357972e-91 | 145 | 2.35017022301052e-99 | 7 |
| CD74.2 | 295 | 1.8192992694437e-83 | 307 | 8.58008635919608e-67 | 6 | 0 | 8 |
| ALDH2 | 112 | 1.73857771420774e-142 | 373 | 6.06047671285392e-54 | 130 | 4.04024967713762e-107 | 9.5 |
| CPVL | 249 | 5.11325442579919e-94 | 132 | 2.02326222241065e-114 | 234 | 1.8552740816104e-77 | 9.5 |

Showing 1 to 10 of 2,314 entries

Previous 1 2 3 4 5 … 232 Next

Download Ranking Consensus

***S6 Fig. A)** Venn diagram showing the overlap of DEGs obtained by the scRNA-Seq pipeline for the 2700 PBMCs data set. **B)** An interactive rank-based consensus table showing the Top 10 highest ranked genes generated with the same data. The diagram shows a considerable agreement between DEGs identified by the three DE analysis methods and a relatively large number of DEGs by both Wilcoxon and MAST.*
